# Temporal dynamics of neuroplasticity and neurodegeneration in the central auditory system following noise-induced hearing loss: A multimodal imaging and histological study

**DOI:** 10.1101/2025.09.10.675281

**Authors:** Víctor Giménez-Esbrí, Susanne Schwitzer, Susanne Mueller, Stefan Koch, Marco Foddis, Stefan Donat, Jan Schmoranzer, Niclas Gimber, Janina Zeqiraj, Dietmar Basta, Philipp Boehm-Sturm, Moritz Gröschel

## Abstract

Noise-induced hearing loss (NIHL) is a sensorineural disorder that triggers profound neuroplastic and neurodegenerative consequences in the central auditory nervous system (CNS). This study examined how neuronal density, axonal integrity and glutamatergic and GABAergic neurotransmission are affected after acute overstimulation in the central inferior colliculus (CIC) and the ventral medial geniculate body of the thalamus (MGV). To achieve this, a correlative multimodal approach combining audiometric, magnetic resonance imaging (MRI) and histological biomarkers was performed. Adult mice were noise-exposed to broadband white noise (5-20 kHz) for 3 hours at either high (115 dB SPL) or moderate (90 dB SPL) intensity, while unexposed mice were used as controls. Separate cohorts of mice were investigated 1-, 7-, 56– and 84-days post-exposure using in vivo magnetic resonance imaging (MRI) techniques: Voxel-based morphometry (VBM) of gray matter density (GMD), diffusion MRI (dMRI) of microstructure and connectivity, and single-voxel proton magnetic resonance spectroscopy (1H-MRS) for glutamate and GABA quantification. Frequency-specific auditory brainstem responses (ABR) were recorded at 4, 8, 16 and 32 kHz before and after exposure to examine hearing threshold (HT) shifts. Following imaging, brains were processed for fluorescence immunohistochemistry (FIHC) targeting Neuronal Nuclear Protein (NeuN), 4’,6-diamidino-2-fenilindo (DAPI), Neurofilament (SMI312), vesicular GABA transporter (VGAT), and vesicular glutamate transporters 1 (VGLUT1) and 2 (VGLUT2). HTs were significantly elevated 7d, 56d and 84d after 115 dB noise exposure, suggesting a NIHL phenotype. Neurofilament density significantly increased in the CIC and MGV 1d after 115 dB noise exposure, and returned to baseline at later time points. dMRI transient microstructural alterations were observed 7d after 90 dB noise exposure. Glutamate and GABA decreases 84d after 90 dB exposure were also detected. Moreover, correlation analysis between audiometric, histological and MRI datasets revealed scarce relationships between the investigated parameters. These findings advance our understanding of the CNS adaptations of NIHL and emphasizes the need for more sensitive and integrative approaches to identify robust biomarkers of central auditory dysfunction.

## INTRODUCTION

Noise-induced hearing loss (NIHL) is defined as the sensorineural deafness resulting from prolonged and repetitive exposure to loud noise (**1**). NIHL has been estimated to affect over 5% of the worldwide population, being the second most common cause of acquired hearing loss after age-related hearing loss (ARHL) (**2–5**). As such, NIHL represents a significant hearing concern with substantial impact on individuals’ quality of life, including increased social stress, social isolation and relationship difficulties, a higher risk of depression, and/or cognitive decline commonly associated with hearing deficits (**4,6,7**).

NIHL is a multifactorial and complex disorder caused by the interaction of genetic and environmental factors, but is primarily determined by the degree of biological injury caused by noise exposure (8). In the peripheral auditory system, multiple biological mechanisms have been proposed in order to explain the wide range of noise-induced alterations, such as death of the sensory Hair Cells (HC), Auditory Nerve Fibers (ANF) and/or Spiral Ganglion Neurons (SGN) (**9,10**); The induction of acute inflammatory responses and increases in reactive oxidative species (ROS) formation (**11,12**); mechanical damage to the entire organ of Corti (OC) (**1,3,13**); and the disruption of calcium homeostasis and metabolic pathways implicated in synaptic transmission, leading to the activation of apoptotic cell death pathways (**12,14**). Following intense, repetitive, and prolonged exposure to loud noise, hearing damage could become permanent and irreversible, primarily due to the inability of mammalian HCs to regenerate over-time (15,16). However, none of the aforementioned peripheral consequences fully explains the pathogenesis of NIHL by its own, which make clinical treatment and preventive strategies for people suffering from NIHL notably challenging (**3,9,17**). Besides the impairment of the peripheral auditory system, key detrimental damage can also affect the entire ascending auditory central nervous system (ACNS) (**18–24**). Thus, noise-induced alterations within the central nervous system (CNS) have been linked to several NIHL-derived pathological mechanisms and/or clinical symptoms, as well as to related hearing disorders such as tinnitus or hyperacusis (**25–29**). These central effects have been increasingly gaining attention in the field due to their essential clinical implications in the NIHL pathology. Therefore, there is increasing evidence suggesting that the ACNS could also have a key role in the generation of the whole NIHL adverse effects.

In the ACNS, crucial neurodegenerative and neuroplastic consequences have been observed after NIHL, which can be summarized in: (**1**) Activation of apoptotic and cell death pathways leading to neuronal and cell loss (**20,30,31**); (**2**) Adaptative compensatory neuroplasticity and tonotopic synaptic reorganizations in order to compensate peripheral auditory deprivation (**22–24,32–36**); and (**3**) Imbalances in the excitatory and inhibitory activity as a consequence of the disruption of glutamatergic and GABAergic synaptic neurotransmission (**32,37–39**). Nevertheless, there are still unsolved arduous challenges in this field. First, how these central effects are produced and how they could concretely lead to pathological consequences is still largely unknown. Moreover, there is little knowledge about how this neurodegeneration and neuroplasticity could fluctuate in a time-course manner after NIHL. Increasing the understanding of which short– and long-term biological mechanisms occur in the CNS after loud noise exposure, will help to develop novel preventive, diagnostic and therapeutic strategies against NIHL. Second, potent therapeutic and diagnostic strategies against NIHL are scarce and notably challenging, especially for central noise-induced consequences (**17**). Although some remarkable advances in research and development on new drugs has been done, and some strategies could be applied in the clinics to alleviate its symptoms (**40**), there are no direct clinically available treatments to deal with this hearing disorder. Finally, there is a lack of effective diagnostic methods to detect NIHL consequences in its early stages, where there is a therapeutic window in which most of the promising treatment strategies could be especially effective.

In this context, advanced imaging techniques have been purposed as a potential clinical strategy to monitor and detect central noise exposure consequences. Magnetic Resonance Imaging (MRI) allows noninvasive assessment brain structure and function, offering the advantage of performing multiple versatile sequences within a single imaging session. The resulting signals have been associated and/or correlated with different aspects of the neurobiology of hearing loss, such as changes in the volume of grey matter density (GMD) or white matter density (WMD) in auditory and non-auditory brain areas of human patients suffering from tinnitus, age-related hearing loss (ARHL) or occupational NIHL (**41–53**). However, the potential of MRI to detect NIHL during its early or long-term stages has yet to be fully elucidated. Similarly, it is not clear how well-described audiometric and histological alterations after noise exposure could translate into potential MRI-derived biomarkers for NIHL.

In the present experiments, we investigated changes in cell density, axonal integrity, and glutamatergic and GABAergic neurotransmission as a consequence of loud noise exposure in two core areas of the ACNS, the Central Inferior Colliculus (CIC) and the Ventral Medial Geniculate Body of the thalamus (MGV). These regions were selected due to their key role in the processing of ascending auditory neural signals and its implications in NIHL pathology. These alterations were evaluated 1 day (1d), 7 days (7d), 56 days (56d) and 84 days (84d) in both ACNS regions of interest in a mouse model of NIHL. It was the aim of this exploratory study to identify the temporal dynamics of neuroplasticity and neurodegeneration in the ACNS after NIHL, as well as to identify noninvasive biomarkers as a correlate for noise-induced pathophysiology, elucidating their potential use as a novel diagnostic tool.

## RESULTS

### PERMANENT THRESHOLD SHIFT AFTER HIGH NOISE EXPOSURE

Auditory threshold shifts were significantly elevated after 7 days post-exposure in the 115 dB group (n = 8) when compared to both 90 dB (p ≤ 0.001 for all tested frequencies) and Ctrl groups (p ≤ 0.001 for all tested frequencies). Furthermore, elevated threshold shifts in the 115 dB group were also observed 56 days and 84 days post-exposure when compared to the 90 dB and Ctrl groups (p ≤ 0.05 and/or p ≤ 0.001 for all tested frequencies). The threshold shifts within the investigated frequencies ranged between 50– and 70-dB SPL in the 115 dB group, whereas a smaller threshold shift between 5– and 10-dB SPL could be observed in the 90 dB group for all investigated time points. Moreover, no significant differences were observed between the 90dB and Ctrl groups within all tested time points (p > 0.05 for all tested frequencies). An even slight increase in auditory threshold shift was observed for the 90 dB group 7 days post-exposure (see **Figure 2**, **Table 1**).

**Figure 1.**
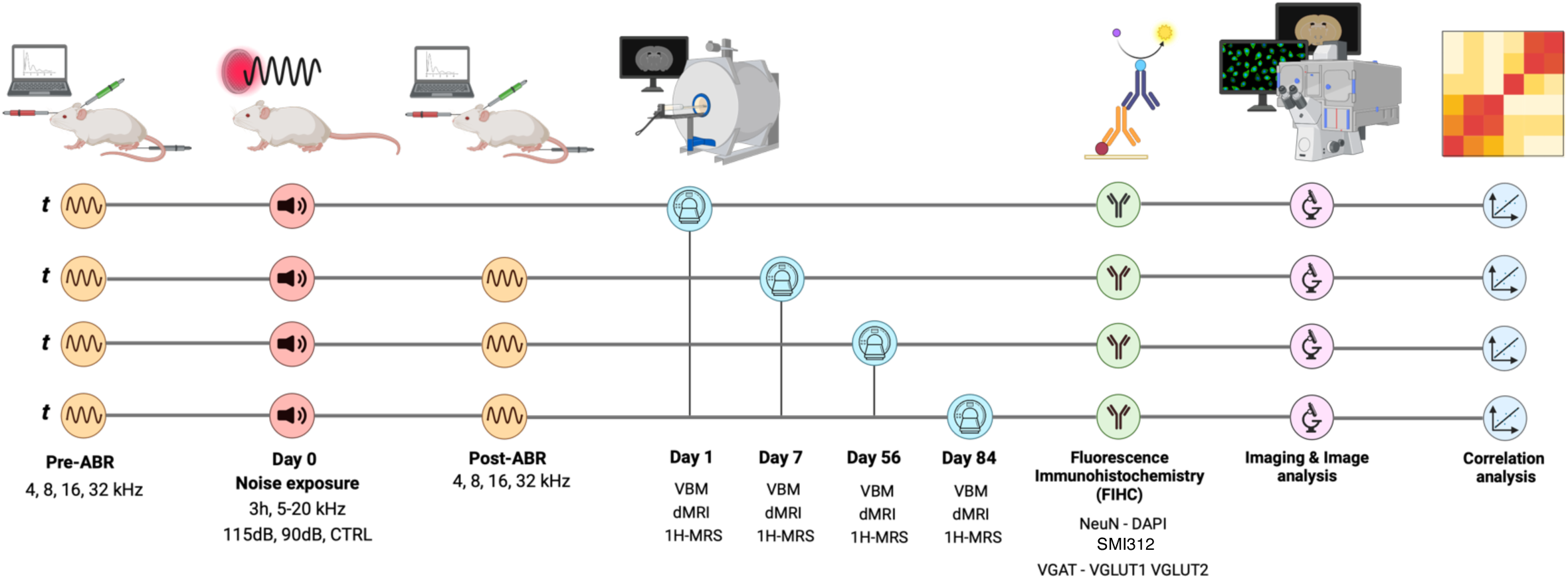
Study Design. NMRI mice were exposed to broadband noise (5–20 kHz, 3 h) at either 115 dB (high exposure) or 90 dB (moderate exposure). Non-exposed animals were used as the control group. Auditory Brainstem Responses (ABR) were acquired before noise exposure (pre-ABR) to assess Hearing Thresholds (HT) of the animals. At 1-, 7-, 56-, and 84-days post-exposure, a second ABR was performed (post-ABR), and MRI scans (VBM, dMRI, and 1H-MRS) were acquired in separate cohorts of mice. Afterwards, brains were then processed for fluorescence immunohistochemistry (FIHC) targeting NeuN, DAPI, SMI312, VGLUT1, VGLUT2, and VGAT. Histological and MRI data were analyzed using both automated and manual methods, followed by correlation analyses to identify relationships between audiometric, MRI and histological markers.

**Figure 2.**
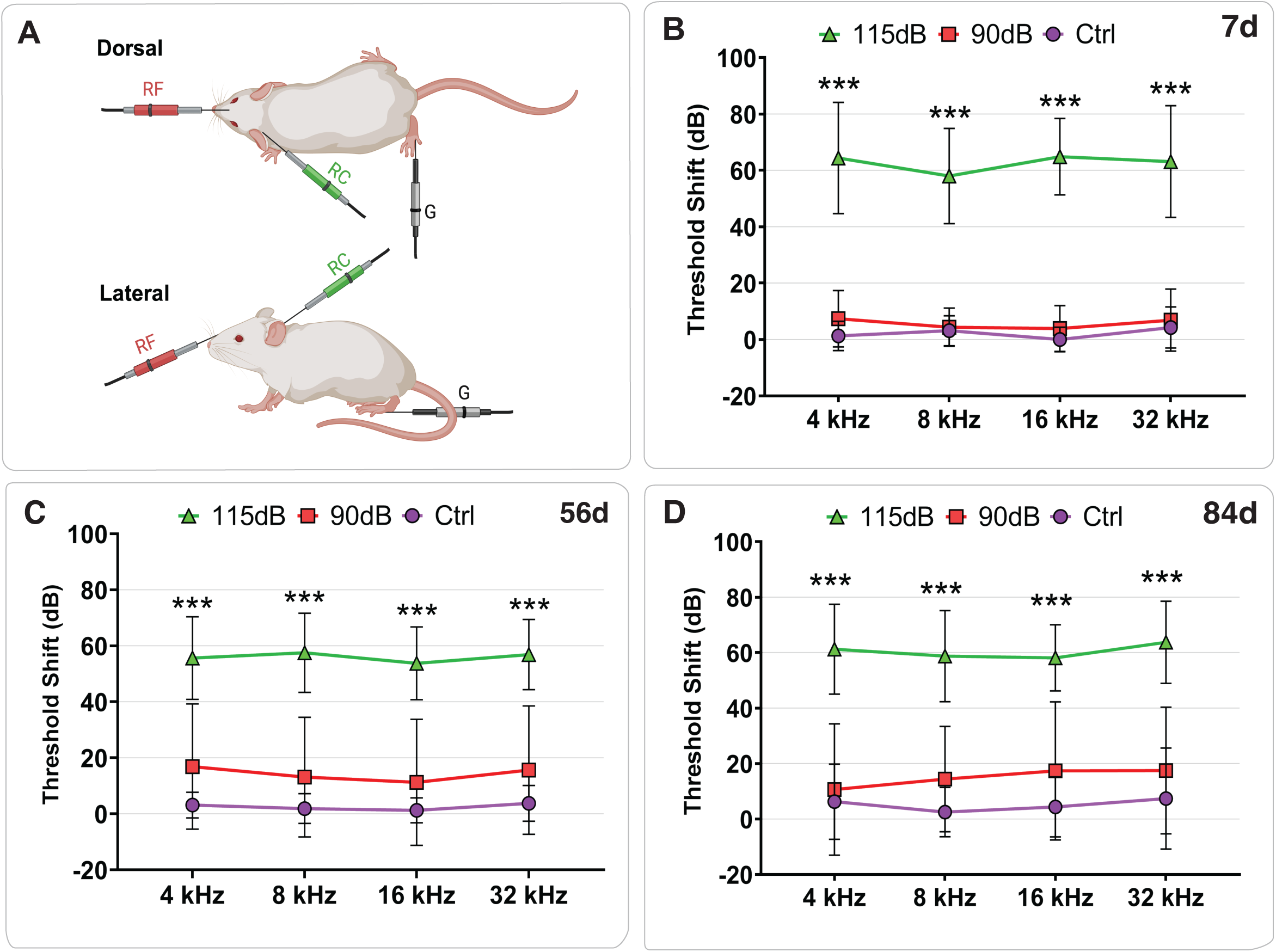
Elevations in Auditory Hearing Threshold after noise exposure. (A) Schematic representation of the Auditory Brainstem Response (ABR) recording setup in mice. Subdermal needle electrodes were placed at the forehead (reference), mastoid (recording), and foot (ground) while animals were anesthetized and housed in a sound-attenuated chamber. Acoustic stimuli were delivered binaurally at four frequencies (4, 8, 16, and 32 kHz), and ABRs were recorded both before (pre-ABR) and after (post-ABR) noise exposure to determine hearing threshold shifts. (B–D) Post-exposure hearing threshold shifts (Mean ± SD) measured at 7 days (B), 56 days (C) and 84 days (D) after noise exposure. Mice exposed to 115 dB showed significantly elevated threshold shifts (50–70 dB) across all frequencies and time points compared to both 90 dB and control groups (p ≤ 0.001). Mice exposed to 90 dB exhibited minimal shifts (5–10 dB), with no significant differences from controls at any time point (p > 0.05). No significant differences were observed between 115 dB groups across time points, nor among control groups (p > 0.05: n.s.; *: p < 0.05; **: p < 0.01, ***: p < 0.001).

**Table 1.** Hearing Threshold shift 7d. 56d and 84d after different noise exposure conditions (Mean ± SD). * p ≤ 0.05; **: p < 0.01; ***: p < 0.001.

| Day | Noise Exposure (dB SPL) | N | Frequency (kHz) |  |  |  |
| --- | --- | --- | --- | --- | --- | --- |
|  |  |  | 4 | 8 | 16 | 32 |
| 7d | 115 dB | 8 | 66.87 ± 21.70 *** | 59.87 ± 19.67 *** | 64.25 ± 13.42 *** | 63.75 ± 20.31 *** |
| | 90 dB | 8 | 6.75 ± 9.57 | 4.33 ± 6.78 | 3.25 ± 7.75 | 5.62 ± 9.79 |
| | 0 dB | 8 | 2.50 ± 4.62 | 5.62 ± 5.62 | 1.12 ± 4.32 | 4.25 ± 7.28 |
| 56d | 115 dB | 8 | 52.00 ± 12.81 *** | 55.62 ± 13.99 *** | 53.12 ± 13.07 *** | 54.37 ± 11.47 *** |
| | 90 dB | 8 | 7.50 ± 6.54 | 6.25 ± 4.43 | 3.75 ± 5.82 | 6.87 ± 5.93 |
| | 0 dB | 8 | 3.57 ± 4.75 | 2.14 ± 5.66 | 1.43 ± 4.75 | 2.85 ± 6.36 |
| 84 | 115 dB | 8 | 61.25 ± 16.20 *** | 58.75 ± 16.42 *** | 58.12 ± 11.93 *** | 63.75 ± 14.82 *** |
| | 90dB | 8 | 6.25 ± 17.47 | 10.00 ± 14.39 | 8.12 ± 13.07 | 8.12 ± 14.37 |
| | 0 dB | 8 | 8.75 ± 11.87 | 4.37 ± 8.63 | 7.50 ± 10.00 | 11.75 ± 16.61 |

### GRAY MATTER DENSITY AND FRACTIONAL ANISOTROPY AFTER NOISE EXPOSURE

No significant changes in GMD were observed between groups either in the CIC or the MGV across the timepoints investigated (p > 0.05 in all tested comparisons by Tukey Post-hoc Test) (see **Figure 3**, **Table 2**). Therefore, a reduction in gray matter was not observed after different noise exposure conditions in the current study. Similarly, Fractional Anisotropy (FA), a marker of microstructural integrity, showed no significant differences between noise exposure conditions within each timepoint (see **Figure 3**, **Table 2**).

**Figure 3.**
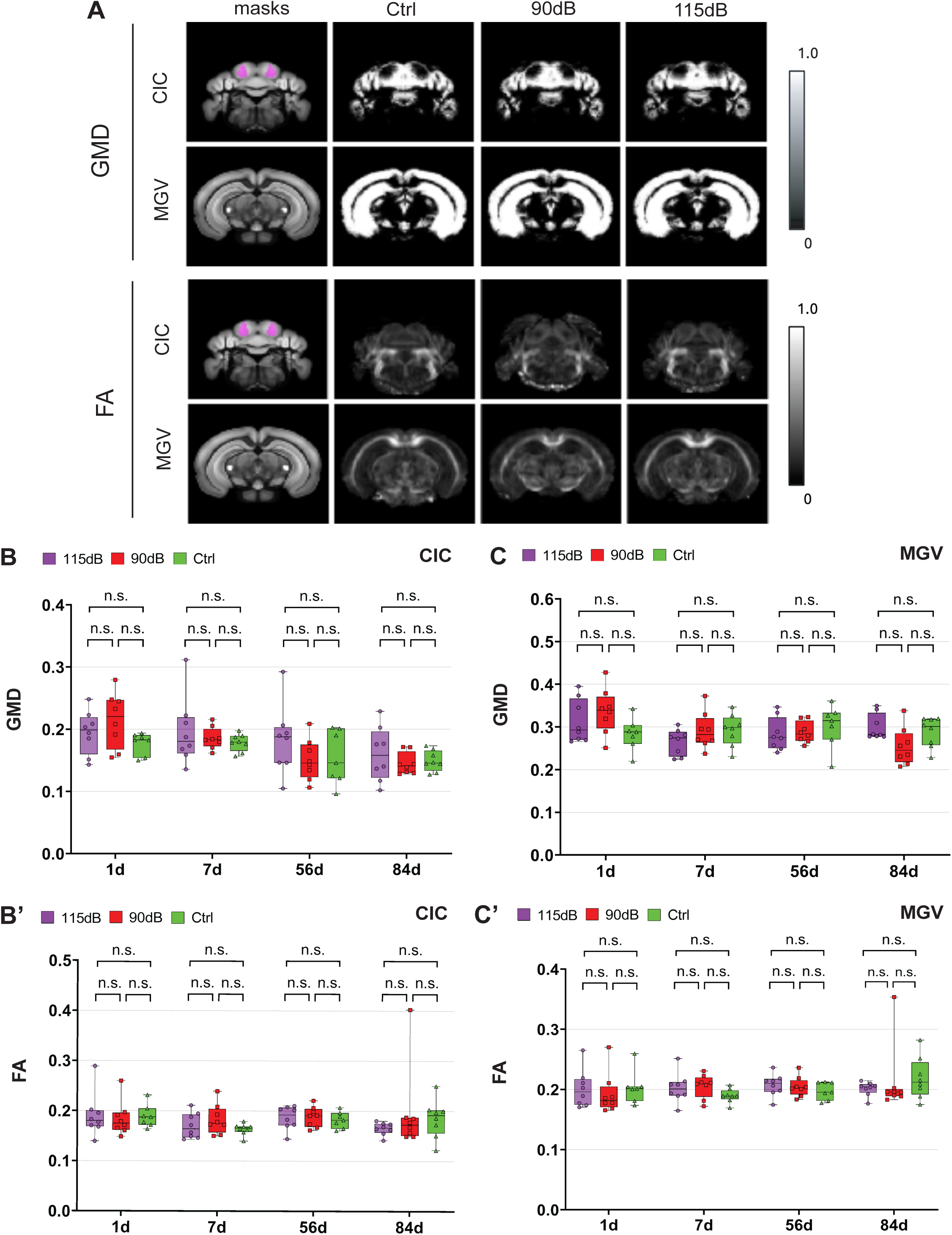
Gray matter density and FA are not significantly altered after noise exposure. **(A)** Representative T2-weighted images showing regions of interest (ROIs) in the CIC and the MGV, as defined using a custom atlas based on the Allen Mouse Brain Atlas. **(B, B′)** Quantification of mean gray matter density (GMD) **(B)** and fractional anisotropy (FA) **(B′)** in the CIC across experimental groups (Ctrl, 90 dB, 115 dB) and time points (1d, 7d, 56d, 84d). **(C, C′)** Quantification of mean gray matter density (GMD) **(C)** and fractional anisotropy (FA) **(C′)** in the MGV across experimental groups (Ctrl, 90 dB, 115 dB) and time points (1d, 7d, 56d, 84d). No significant changes in GMD or FA were observed in either region at any time point or exposure condition (p > 0.05: n.s.; *: p < 0.05; **: p < 0.01, ***: p < 0.001).

**Table 2.** Gray Matter Density (GMD), Fractional Anisotropy (FA), and DAPI^+^ and NeuN/DAPI^+^ cell counts after noise exposure in the CIC and the MGV (Mean ± SD). * p ≤ 0.05; **: p < 0.01; ***: p < 0.001.

| Day | Noise Exposure (dB SPL) | N | CIC |  |  |  | MGv |  |  |  |
| --- | --- | --- | --- | --- | --- | --- | --- | --- | --- | --- |
|  |  |  | GMD | FA | DAPI+ | NeuN+/DAPI+ | GMD | FA | DAPI+ | NeuN+/DAPI+ |
| 1d | 115dB | 8 | 0.19 ± 0.03 | 0.19 ± 0.04 | 1171.61 ± 149.48 | 903.12 ± 156.04 | 0.31 ± 0.05 | 0.20 ± 0.03 | 750.63 ± 54.54 | 493.01 ± 46.44 |
| | 90dB | 8 | 0.21 ± 0.04 | 0.18 ± 0.03 | 1143.97 ± 91.40 | 827.29 ± 159.00 | 0.33 ± 0.05 | 0.19 ± 0.03 | 745.33 ± 28.70 | 487.20 ± 70.08 |
| | 0 dB | 7 | 0.177 ± 0.01 | 0.18 ± 0.02 | 1092 ± 178.60 | 821.87 ± 230.86 | 0.28 ± 0.03 | 0.20 ± 0.02 | 714.45 ± 73.93 | 458.46 ± 71.43 |
| 7d | 115 dB | 8 | 0.19 ± 0.05 | 0.17 ± 0.02 | 1361.50 ± 109.24 | 1102.48 ± 100.98 | 0.26 ± 0.03 | 0.20 ± 0.02 | 863.74 ± 104.38 | 564.16 ± 86.20 |
| | 90 dB | 8 | 0.18 ± 0.01 | 0.18 ± 0.02 | 1346.61 ± 95.42 | 1104.98 ± 130.56 | 0.29 ± 0.04 | 0.20 ± 0.19 | 839.49 ± 57.20 | 580.54 ± 55.90 |
| | 0 dB | 8 | 0.17 ± 0.01 | 0.16 ± 0.01 | 1238.20 ± 135.10 | 965.94 ± 207.73 | 0.29 ± 0.03 | 0.18 ± 0.11 | 821.23 ± 77.62 | 543.70 ± 98.25 |
| 56d | 115 dB | 8 | 0.18 ± 0.55 | 0.18 ± 0.02 | 1244.94 ± 192.04 | 974.58 ± 184.22 | 0.28 ± 0.03 | 0.20 ± 0.01 | 791.94 ± 56.27 | 538.64 ± 56.99 |
| | 90 dB | 8 | 0.15 ± 0.03 | 0.18 ± 0.02 | 1208.67 ± 164.47 | 937.45 ± 207.71 | 0.28 ± 0.02 | 0.20 ± 0.01 | 766.27 ± 94.03 | 487.28 ± 70.51 |
| | 0 dB | 7 | 0.15 ± 0.04 | 0.18 ± 0.01 | 1163.23 ± 164.47 | 924.23 ± 220.97 | 0.30 ± 0.05 | 0.19 ± 0.01 | 758.91 ± 67.65 | 536.64 ± 56.99 |
| **84d** | **115 dB** | 8 | 0.16 ± | 0.16 ± | 1255.16 ± | 837.47± | 0.29 ± | 0.20 ± | 774.13 ± | 495.66 ± |
|  |  |  | 0.04 | 0.01 | 194.99 | 185.63 | 0.03 | 0.01 | 127.45 | 136.97 |
|  | **90dB** | 8 | 0.14 ± | 0.19 ± | 1197.13 ± | 900.90 ± | 0.25 ± | 0.21 ± | 770.14 ± | 510.72 |
|  |  |  | 0.01 | 0.08 | 171.23 | 180.38 | 0.04 | 0.05 | 50.04 | ± 53.69 |
|  | **0 dB** | 8 | 0.14 ± | 0.18 ± | 121.15 ± | 914.47 ± | 0.28 ± | 0.21 ± | 712.95 ± | 434.65 ± |
|  |  |  | 0.01 | 0.03 | 68.13 | 169.59 | 0.03 | 0.03 | 88.01 | 109.87 |

### CHANGES IN NEUN^+^/DAPI^+^ AND DAPI^+^ AND DENSITY IN THE CIC AND MGV

In both areas investigated, significant differences in NeuN^+^/DAPI^+^ cell counts were not detected between the 115 dB group when compared either to the 90 dB or the Ctrl group after noise exposure across the different timepoints included in the study (p > 0.05 for all tested comparisons) (see **Figure 4**, **Table 2**). In the same line, no significant differences in overall cell density (DAPI^+^) were observed between the 115 dB group to the 90 dB and to the Ctrl groups at any timepoint (p > 0.05 in all tested comparisons by Games-Howell Post-hoc test) (see **Figure 4, Table 2**).

**Figure 4.**
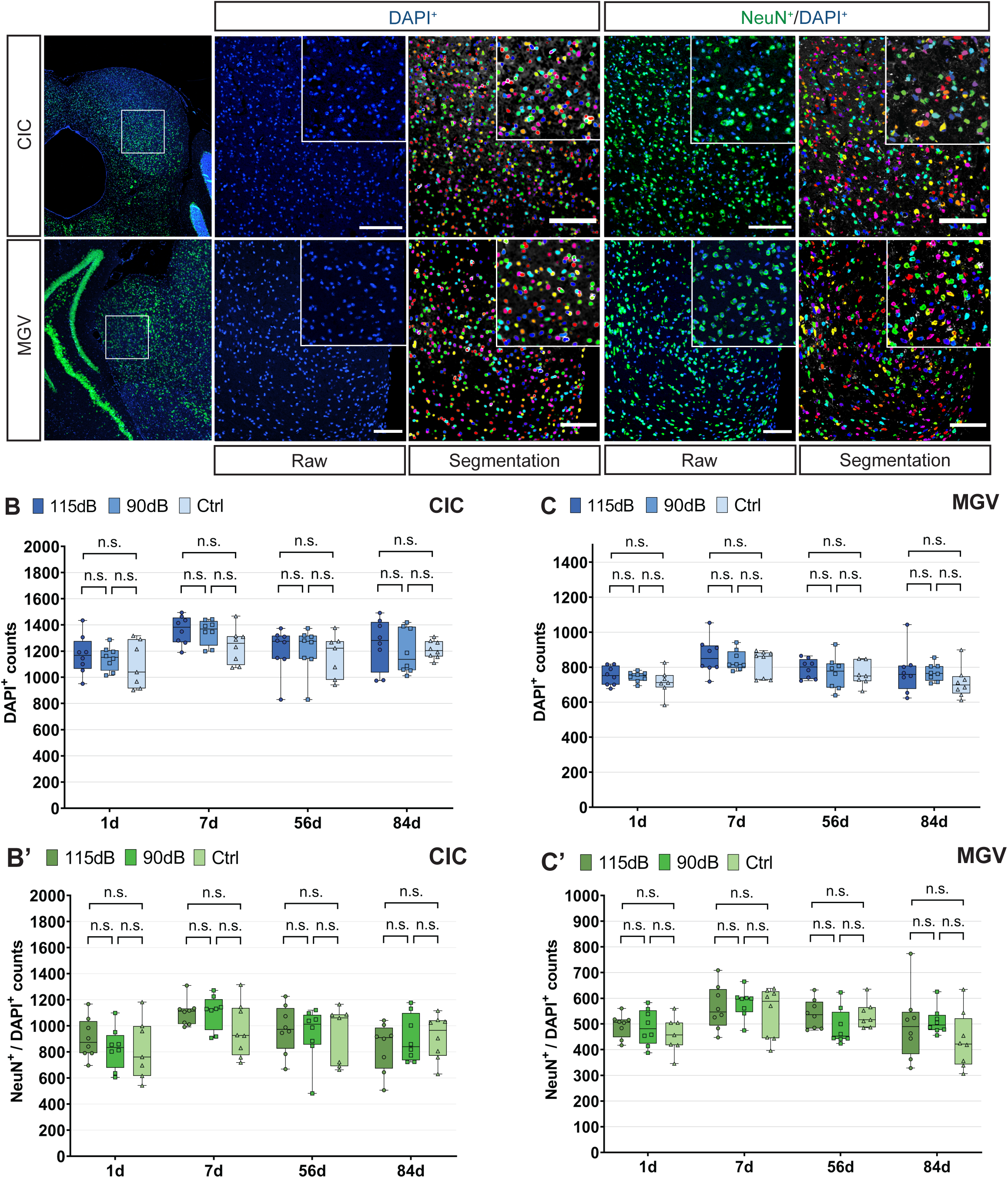
Cell and neuronal density analysis after noise exposure. **(A)** Histological images from Neuronal (NeuN+/DAPI+) and Cell (DAPI+) labeling detected after acute noise exposure in the Central Inferior Colliculus (CIC) **(top)** and Ventral Medial Geniculate Body (MGV) **(bottom)**. Original images and their respective automatic segmentation are shown for both NeuN+/DAPI+ and DAPI+ labeling. Total DAPI+ cell countings in the CIC **(B)** and the MGV **(C)** 1d, 7d, 56d and 84d after noise exposure (115dB: dark blue, 90dB: blue, Ctrl: light blue). No significant (n.s) differences were observed between any experimental treatment either within 1, 7, 56 and 84d after exposure. Total NeuN+/DAPI+ neuronal countings in the CIC **(B’)** and the MGV **(C’)** assessed 1d, 7d, 56d and 84d after noise exposure (115dB: dark green, 90dB: green, Ctrl: light green). No significant (n.s) differences in neuronal density were detected between the experimental groups included in the present study (Mean ± SD, p > 0.05: n.s.; *: p < 0.05; **: p < 0.01, ***: p < 0.001). **Scale bar: 100 μM.**

### WHITE MATTER INTEGRITY CHANGES IN THE CIC AND MGV

Connectivity of CIC and MGV was evaluated via the structural connectome, i.e., streamline reconstructions from the dMRI data. At 7d post-exposure, an increase in streamlines could be observed in the 90 dB group when compared to both 115dB (p = 0.003 by Tukey Post-hoc comparisons) and Ctrl (p = 0.015 by Tukey Post-hoc comparisons) for the MGV (See **Figure 5**, **Table 3**) Those effects were not seen between the 115 dB and Ctrl groups (p > 0.05 by Tukey Post-hoc comparisons), nor for all comparisons performed in the CIC (p > 0.05 by Tukey Post-hoc comparisons). Moreover, no significant differences were found 1d, 56d or 84d post-exposure in either region (p > 0.05 in all tested comparisons by Tukey or Games-Howell Post-hoc Test) (see **Figure 5**, **Table 3**). Likewise, other microstructural integrity indices, including Axial Diffusivity (AD), Radial Diffusivity (RD) and Mean Diffusivity (MD) also showed no significant differences between groups (**Supplemental Fig. 1**).

**Figure 5.**
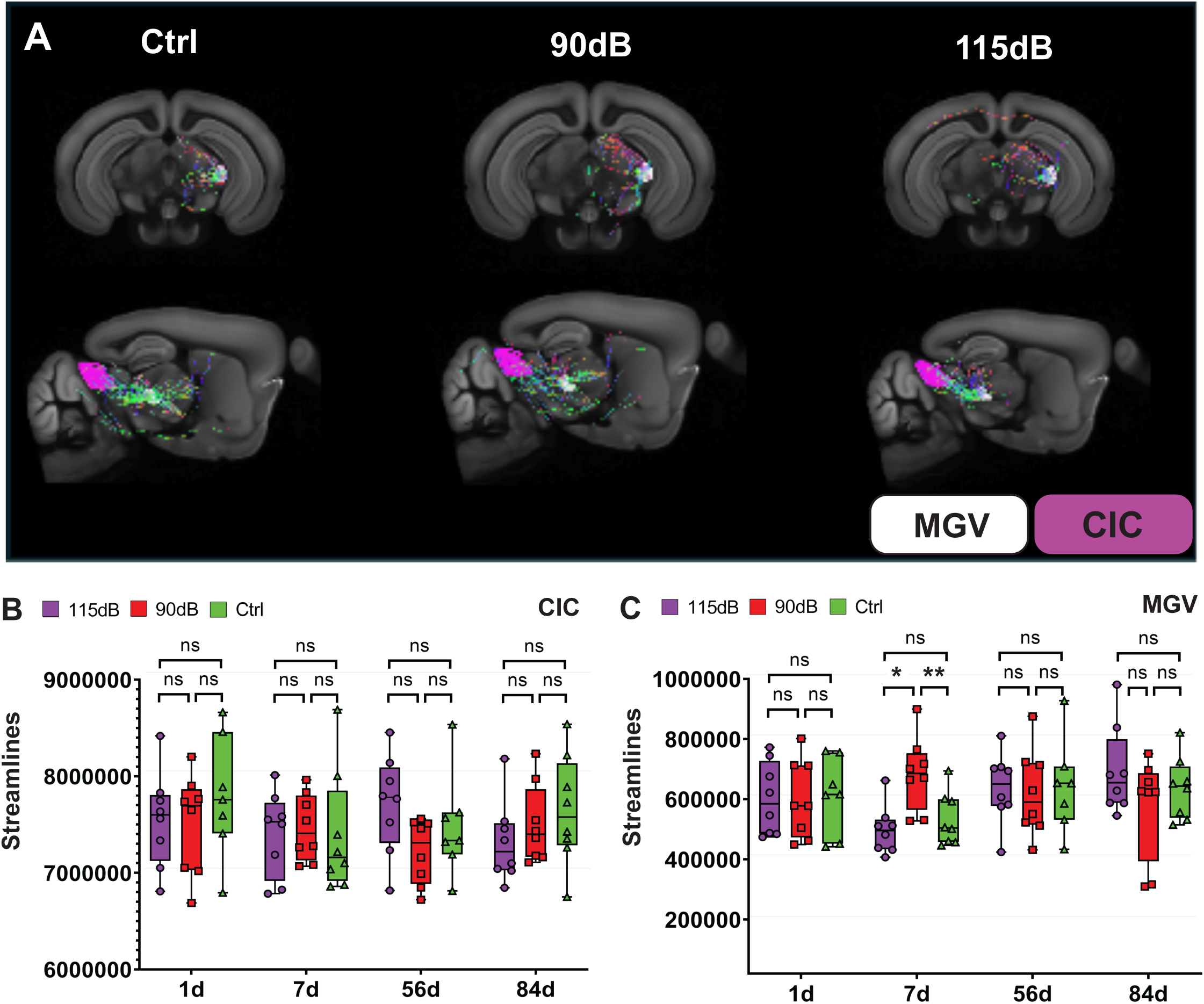
Diffusion MRI (dMRI) connectivity after noise exposure. **(A)** Representative dMRI streamline reconstructions of the fiber tract from CIC (magenta) to MGV (white) in mice 1d after exposure to no (ctrl), 90 dB and 115 dB noise overlaid on Allen brain template. In the CIC, no significant differences (n.s.) in streamline count **(B)** were found in specific tracts (p > 0.05 in all tested comparisons). In the MGV **(C)** significant elevations of streamlines were observed 7d post-exposure between the 90dB and both the Ctrl and 115dB group (p > 0.05: n.s.; *: p < 0.05; **: p < 0.01, ***: p < 0.001).

**Table 3.**
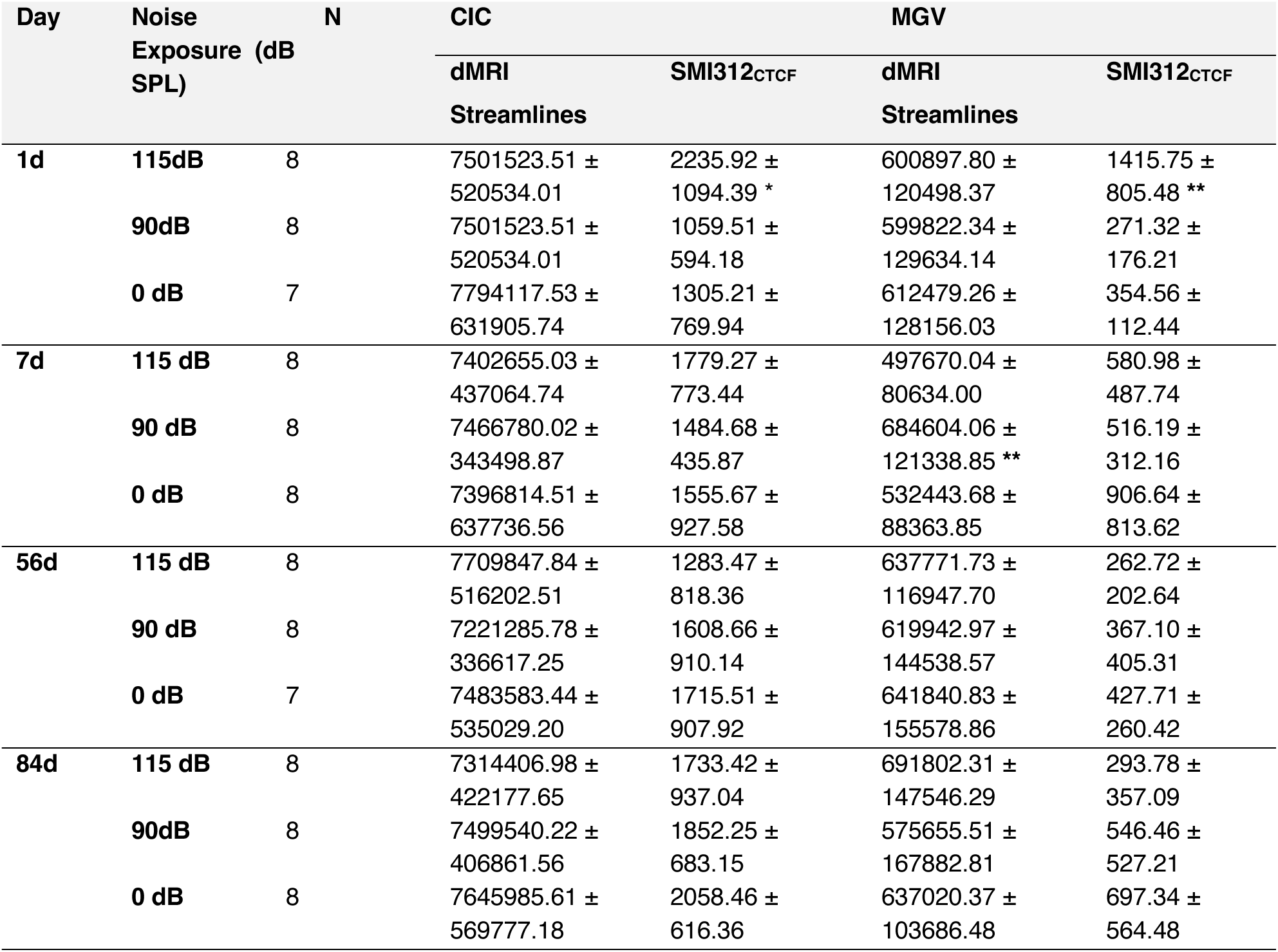
Streamline dMRI reconstruction and SMI312C_TCF_ values after noise exposure in the CIC and the MGV (Mean ± SD). * p ≤ 0.05; **: p < 0.01; ***: p < 0.001.

### EFFECTS OF NIHL ON SMI312 EXPRESSION

In the CIC, a significant increase in SMI312 expression was detected 1d after noise exposure between the 115 dB and the 90dB (p = 0.030 by Tukey Post-hoc test) groups, but no significant differences were found when comparing the 115 dB to the Ctrl group (p = 0.111 by Tukey Post-hoc test). Furthermore, no significant differences were also observed between the 90 dB and Ctrl group (p = 1.000 by Tukey Post-hoc test). 7d, 56d or 84d after noise exposure, no statistically significant differences were found in SMI312 expression between groups (p > 0.05 in all tested comparisons by Tukey Post-hoc Test). However, a temporal decrease of SMI312 expression could be observed from 1d to 84d post-exposure in the CIC (see **Figure 6**, **Table 3**).

**Figure 6.**
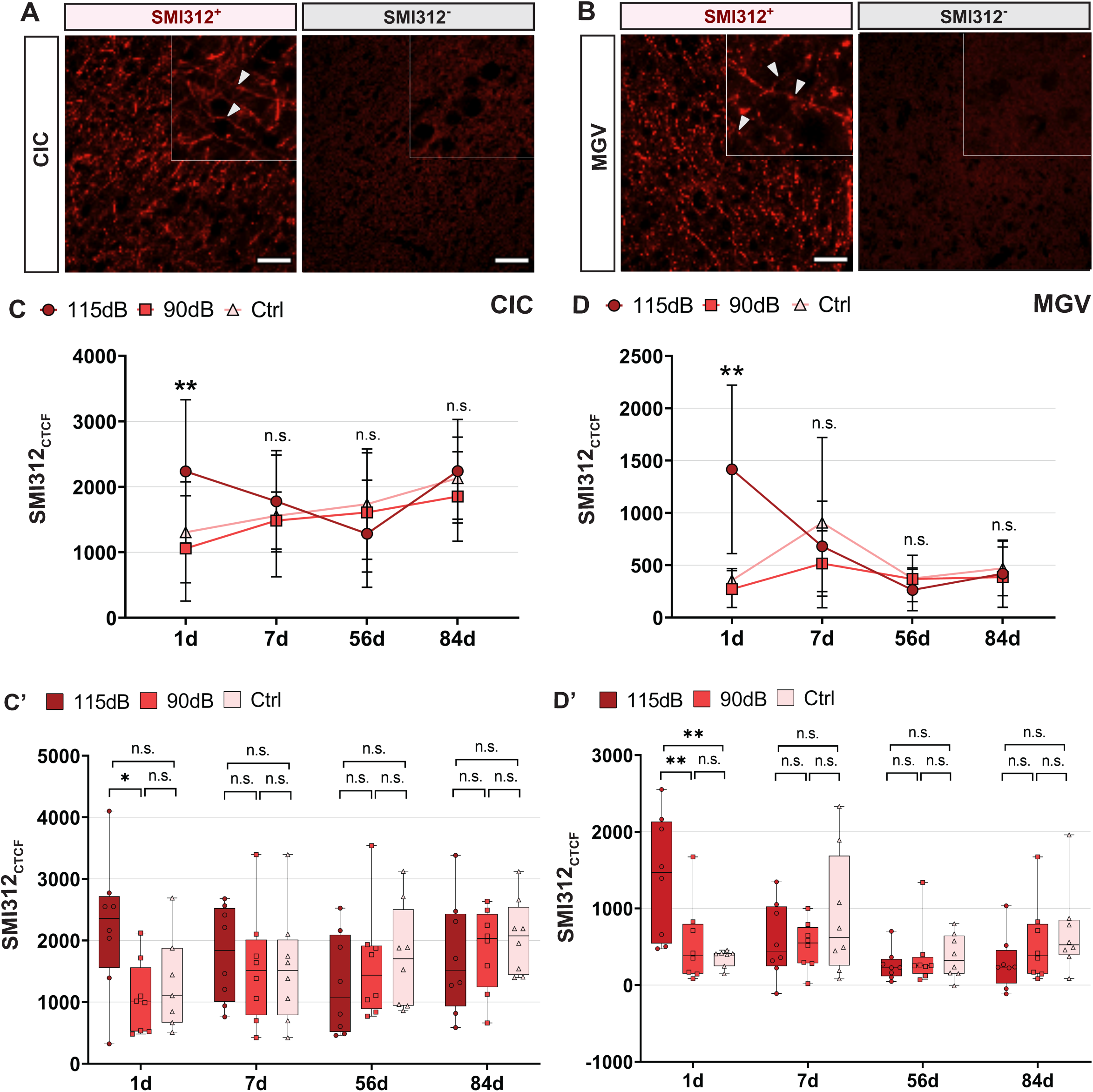
Axonal neuroplastic changes after noise trauma. (A-A’) Representative images showing SMI312 labelling (red) in the CIC **(A)** and the MGV **(A’)**. In both areas, positive (SMI312^+^) and negative (SMI312^-^) labelling could be seen. Arrowheads highlight neurofilaments observed. No specific labelling was observed on SMI312^-^ slices when compared to SMI312^+^ ones. **(C-C’)** Quantitative SMI312 fluorescence intensity measurements (SMI312_CTCF_) in the CIC, showed in barplots **(C)** and boxplots **(C’)**. Significant increases in SMI312_CTCF_ were observed in the CIC 1d post-exposure between the 115 dB and the 90dB groups, whereas no significant (n.s.) differences were detected between the 115 dB and the Ctrl groups. No additional differences were found between the 90 dB and the and Ctrl groups. Moreover, a transient decline of SMI312 expression is shown from 1d to 84d post-exposure. **(D-D’)** Quantitative SMI312_CTCF_ in the MGV, represented both in barplots **(D)** and boxplots **(D’)**. In the MGV, the 115 dB group showed significantly increased SMI312_CTCF_ levels when compared to both the 90 dB and Ctrl groups. 7d, 56d and 84d after acute noise exposure, no significant differences were found between any experimental treatment in either in the MGV or the CIC (p > 0.05: n.s.; *: p < 0.05; **: p < 0.01, ***: p < 0.001). Finally, time-course reductions in SMI312 expression are also in the MGV. **Scale Bar: 5 μm**.

For the MGV, a significant increase in SMI312 expression was observed in the 115 dB group when compared to both the 90 dB (p = 0.012 by Games-Howell post-hoc test) and Ctrl (p = 0.017 by Games-Howell post-hoc test) groups 1d post-exposure. No significant differences were observed between the 90 dB and Ctrl mice (p = 0.530 by Games-Howell post-hoc test). As it was shown for the CIC, no significant changes in SMI312 expression were observed 7d, 56 or 84d post-exposure between the different noise exposure conditions (p > 0.05 by Tukey post-hoc test). Moreover, the temporal decline in SMI312 expression over-time could also be observed within this region (see **Figure 6**, **Table 3**).

### 1H MRS OF NEUROTRANSMITTER CONCENTRATIONS

Slight Glutamate and GABA concentration changes were found mainly 84 after noise exposure (see **Figure 7**, **Table 4**). At this timepoint, a significant decrease in glutamate concentration was found in the 90dB group when compared to the Ctrl group (p = 0.048 by Tukey Post-hoc test). However, this decrease was not significant between the 90 dB and the 115 dB group (p > 0.05 by Tukey Post-hoc test) or the 115dB and Ctrl groups (p > 0.05 by Tukey Post-hoc test). Regarding GABA, a significant decrease in concentration was also found in the 90 dB group when compared to the Ctrl group (p = 0.048 by Tukey Post-hoc test). When compared to the 115 dB group, although not significant, a trend towards a decrease in GABA concentration was found (p = 0.066 by Tukey Post-hoc test).

**Figure 7.**
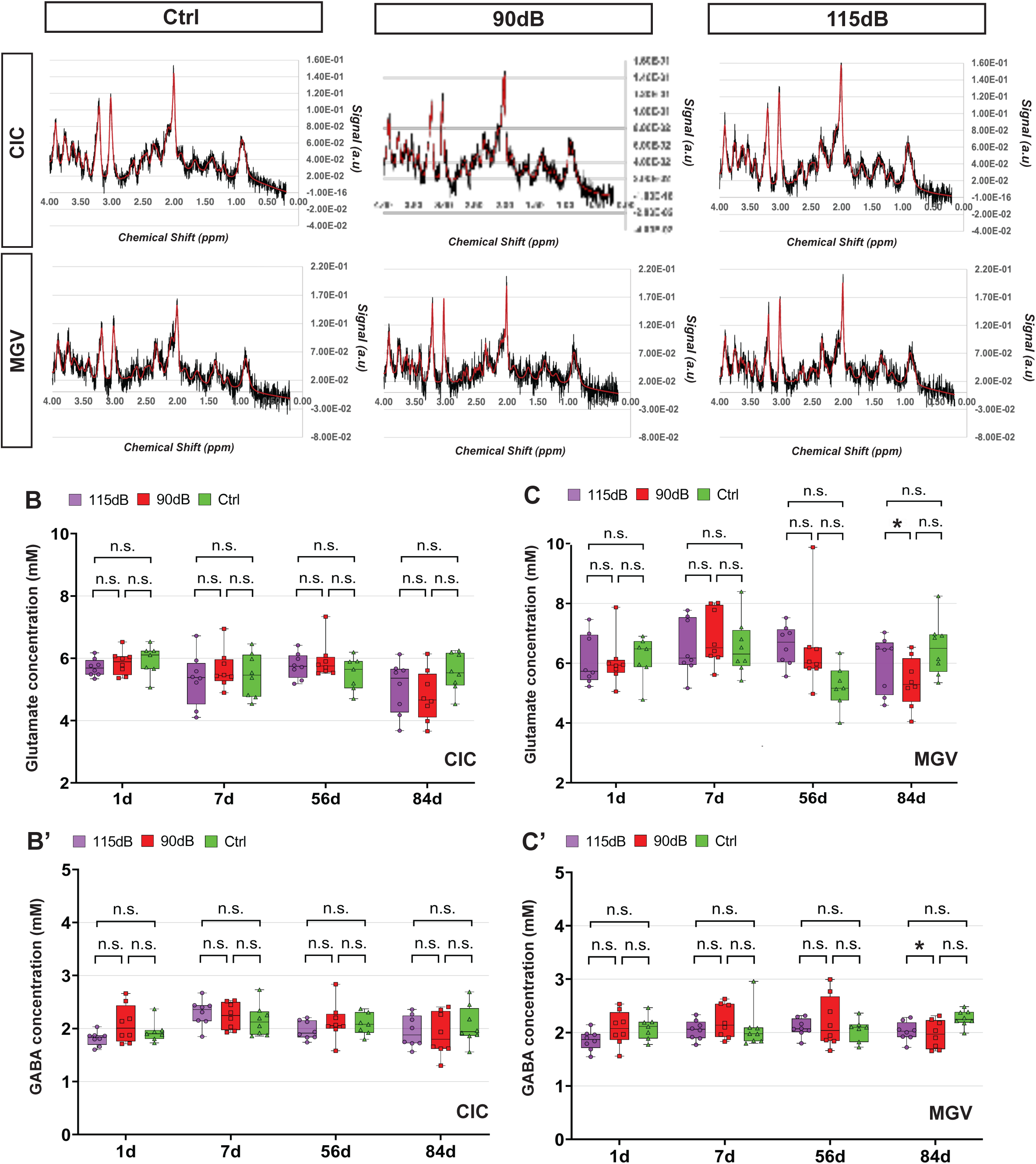
Glutamate and GABA spectroscopy analysis after noise exposure. **(A)** Representative 1H-MRS spectra of the CIC and MGV of mice with no (ctrl), 90 db and 115 db noise trauma. Raw (black) and fitted data (red) are shown, peaks of main metabolites are indicated in the IC/ctrl spectrum **(A)**. Quantification in the CIC for glutamate **(B)** and GABA **(B’)**, and in the MGV for glutamate **(C)**, and GABA **(C’)**. Small decreases in Glutamate and GABA concentrations were found 84d after exposure in the MGV in the 90dB group when compared to Ctrls. No differences were found between 115 dB-exposed mice. No additional significant differences in GABA and Glu were detected in the respective groups. NAA: N-Acetylaspartate; Cr: Creatine; PCr: Phosphocreatine; Glu: Glutamate; Gln: Glutamine; mI: myo-Inositol; GABA: γ-Aminobutyric Acid; GSH: Glutathione; Cho: Choline; NAAG: N-Acetylaspartylglutamate (p > 0.05: n.s.; *: p < 0.05; **: p < 0.01, ***: p < 0.001).

**Table 4.** Glutamate and GABA concentration (nM) values by means of 1H-MRS after noise exposure in the CIC and the MGV (Mean ± SD). * p ≤ 0.05; **: p < 0.01; ***: p < 0.001.

| Day | Noise Exposure (dB SPL) | N | CIC |  | MGV |  |
| --- | --- | --- | --- | --- | --- | --- |
|  |  |  | Glutamate (nM) | GABA (nM) | Glutamate (nM) | GABA (nM) |
| 1d | 115dB | 8 | 5.70 ± 0.28 | 1.18 ± 0.13 | 6.11 ± 0.76 | 1.85 ± 0.18 |
| | 90dB | 8 | 5.85 ± 0.38 | 2.08 ± 0.35 | 6.04 ± 0.80 | 2.08 ± 0.32 |
| | 0 dB | 7 | 5.9 ± 0.49 | 1.94 ± 0.20 | 6.18 ± 0.72 | 2.09 ± 0.22 |
| 7d | 115 dB | 8 | 5.3 ± 0.84 | 2.29 ± 0.24 | 6.52 ± 0.93 | 2.04 ± 0.18 |
| | 90 dB | 8 | 5.7 ± 0.64 | 2.26 ± 0.25 | 6.95 ± 0.96 | 2.21 ± 0.33 |
| | 0 dB | 8 | 5.4 ± 0.7 | 2.17 ± 0.30 | 6.74 ± 0.98 | 2.08 ± 0.37 |
| 56d | 115 dB | 8 | 5.52 ± 0.50 | 2.04 ± 0.21 | 5.28 ± 0.70 | 2.08 ± 0.22 |
| | 90 dB | 8 | 5.93 ± 0.59 | 2.13 ± 0.35 | 6.44 ± 1.46 | 2.19 ± 0.47 |
| | 0 dB | 7 | 5.51 ± 0.54 | 2.08 ± 0.20 | 5.22 ± 0.73 | 2.04 ± 0.21 |
| 84d | 115 dB | 8 | 5.06 ± 0.84 | 1.94 ± 0.29 | 5.98 ± 0.94 | 2.02 ± 0.17 |
| | 90dB | 8 | 4.79 ± 0.82 | 1.89 ± 0.40 | 5.36 ± 0.82 * | 1.96 ± 0.26 * |
| | 0 dB | 8 | 5.56 ± 0.60 | 2.06 ± 0.36 | 6.51 ± 0.93 | 2.26 ± 0.15 |

For the other timepoints investigated, no significant differences were found across noise exposure conditions (p > 0.05 in all tested comparisons by Tukey or Games-Howell Posthoc comparisons).

### CHANGES IN VNTT EXPRESSION

Regarding changes in glutamatergic influences, no significant differences were found in the CIC and the MGV in VGLUT1 and/or VGLUT2 expression between the investigated experimental treatments and timepoints studied (p > 0.05 in all tested comparisons by Tukey Post-hoc test). Thus, no significant changes in glutamatergic neurotransmission could be detected by histological characterization (see **Figure 8**, **Table 5**). With regard to GABAergic activity, no significant changes in VGAT expression were also not observed in the CIC or the MGV after acute noise exposure. (p > 0.05 for all tested comparisons by Tukey Post-hoc test) (see **Figure 8**, **Table 5**). Thus, the present histological experiments did not detect significant changes in glutamatergic and GABAergic VNTT expression.

**Figure 8.**
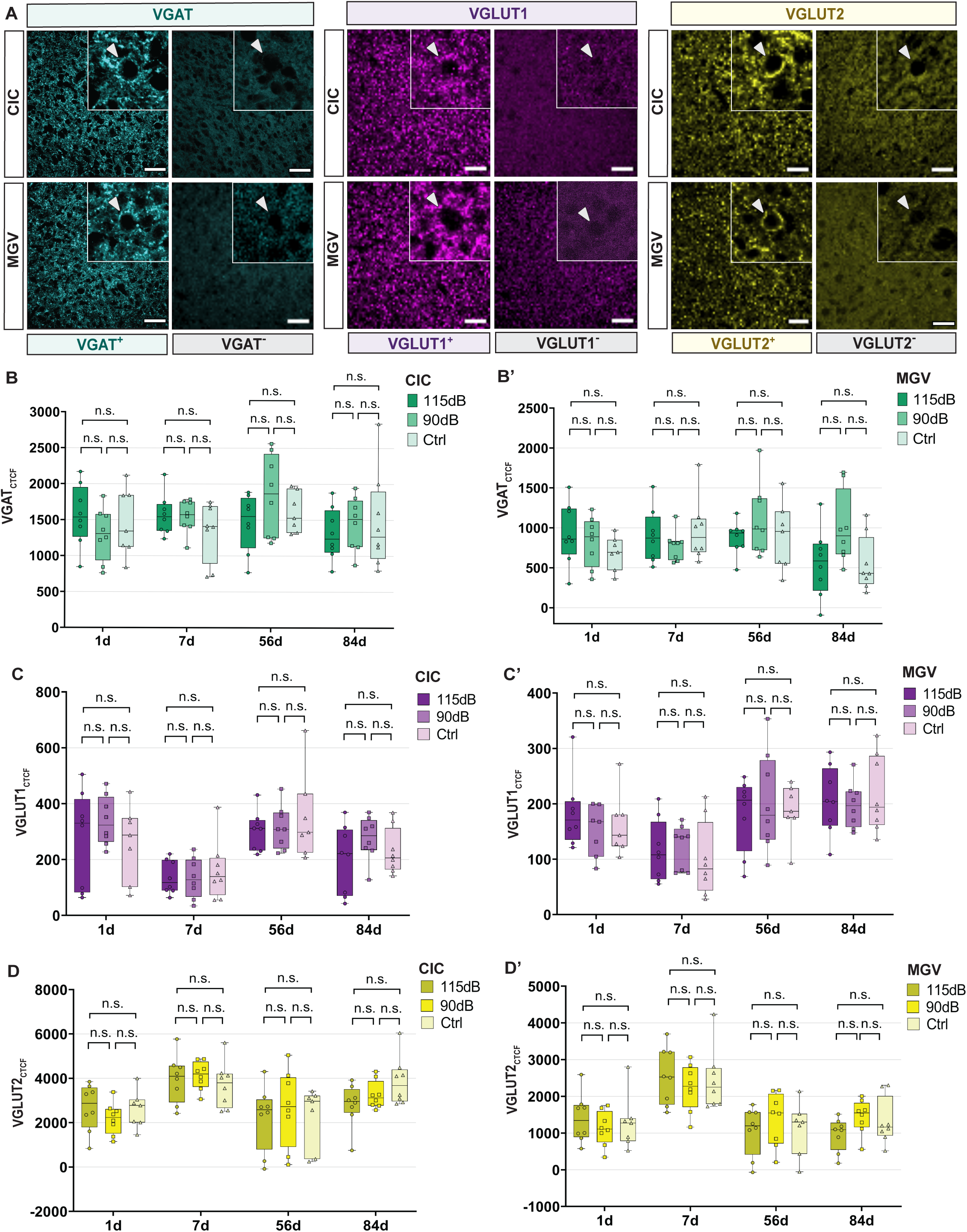
VNTT expression analysis following NIHL. **(A)** Representative images from VGAT (cyan, left), VGLUT1 (magenta, middle) and VGLUT2 (yellow, right) stainings in the CIC (Top) and MGV (Bottom) showing both positive (left) and negative (right) labelling. Arrowheads indicate VNTT staining, surrounding neuronal bodies. No positive labelling was observed on negative control slices surrounding neuronal bodies. Quantitative VGATCTCF **(B-B’)**, VGLUT1CTCF **(C-C’)**, VGLUT2CTCF **(D-D’)** measurements 1-, 7-, 56-, and 84-days post-exposure in the CIC **(left)** and MGV **(right)**. No significant differences were observed in the CIC and MGV for all VNTT measurements between the 115 dB, the 90 dB and/or the Ctrl groups, within any time point investigated. VGAT Look-up Tables (LUTs) were manually adjusted to cyan. VGLUT1 and VGLUT2 are shown in raw LUTs. (p > 0.05: n.s.; *: p < 0.05; **: p < 0.01, ***: p < 0.001). Scale Bar: 5 μm.

**Table 5.**
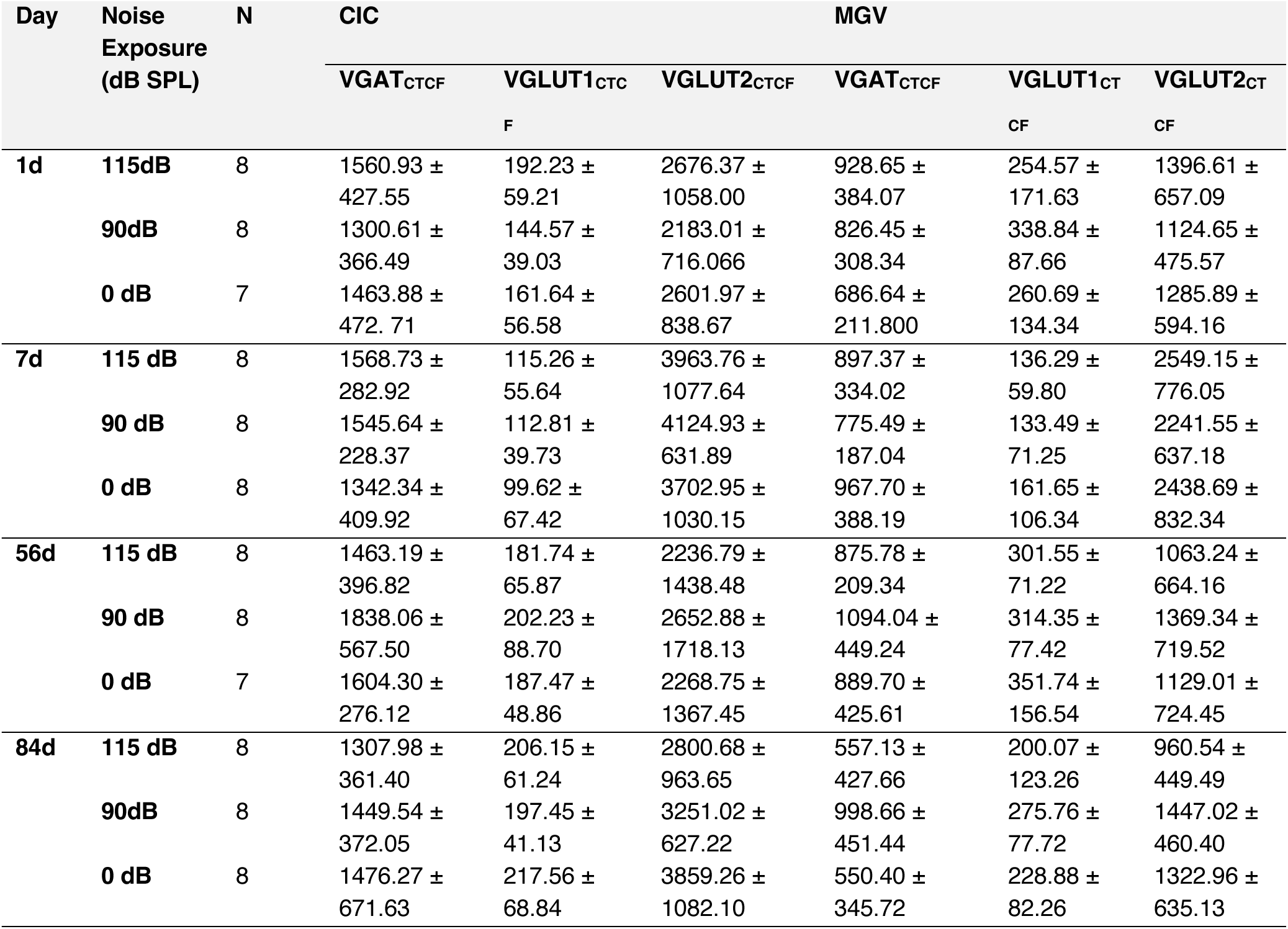
VGAT_CTCF_, VGLUT1_CTCF_ and VGLUT2_CTCF_ values after noise exposure in the CIC and the MGV (Mean ± SD). * p ≤ 0.05; **: p < 0.01; ***: p < 0.001.

### CORRELATION ANALYSIS

To evaluate potential associations between auditory function, MRI-derived parameters, and histological biomarkers, we performed a comprehensive correlation analysis using Spearman’s rank correlation coefficient. Scarce correlation between ABR, MRI and Histological parameters investigated was obtained. The findings are summarized in **Figure 9**.

**Figure 9.**
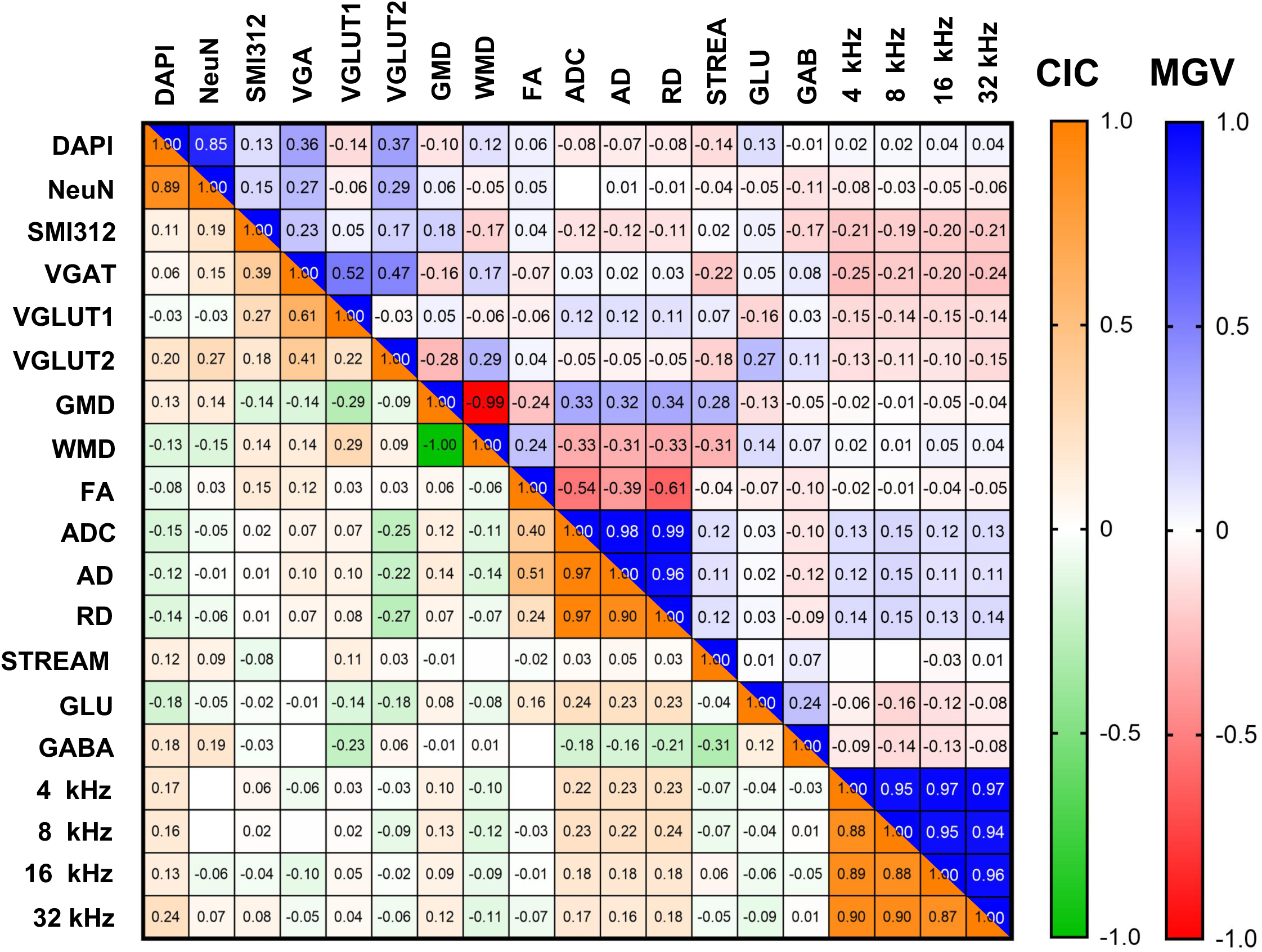
Spearman correlation analysis. Heatmap representation of Spearman correlation coefficients between ABR thresholds, MRI-derived parameters, and histological markers in the Central Inferior Colliculus **(CIC, Upper Triangle)** and Ventral Medial Geniculate Body of the Thalamus **(MGV, Lower Triangle)** Color gradients represent the strength and direction of Spearman correlation coefficient ranks (CIC: +1 = orange, −1 = green; MGV: +1 = blue, −1 = red). Both in the CIC and the MGV, minimal significant correlations were identified between ABR, MRI and FIHC data (*: p < 0.05; **: p < 0.01, ***: p < 0.001; ****: p < 0.00001).

## DISCUSSION

The present study explored the temporal dynamics of neuroplastic and neurodegenerative changes in the central auditory nervous system following acute noise exposure, focusing on neuronal cell density, axonal connectivity and glutamatergic and GABAergic neurotransmission imbalances. Using a multimodal approach combining auditory brainstem response (ABR), magnetic resonance imaging (MRI), and fluorescence immunohistochemistry (FIHC), it was the aim to identify potential early noninvasive biomarkers for noise-induced hearing loss (NIHL). Our findings revealed significant elevations in ABR thresholds in mice exposed to 115 dB, indicating a persistent NIHL phenotype. Increases in SMI312 expression were observed in the CIC and MGV 1d post-exposure followed by a temporal expression decline, suggesting a time-course degeneration of axonal density. Connectivity alterations were also detected by diffusion MRI (dMRI) 7 days after moderate exposure. No significant alterations were detected in neuronal density or glutamatergic and GABAergic markers at any time point investigated. These results highlight early and transient neuroplastic responses in the auditory CNS following noise trauma, while also revealing the challenges in identifying robust noninvasive biomarkers for NIHL.

### Increases in HT and axonal degeneration

The ABR results confirmed that mice exposed to 115 dB experienced significant and persistent threshold shifts across all tested frequencies and time points, consistent with a robust NIHL phenotype and prior reports of peripheral auditory damage conditions (**18,19,36**). In contrast, 90 dB exposure did not lead to permanent hearing loss, in line with previous findings indicating that moderate noise levels induce only transient deficits (**10,54**). Those increases in HTs were accompanied by a marked increase in SMI312 expression in the MGV 1d after exposure to 115 dB noise, with a similar but less pronounced change in the CIC. A later decline in SMI312 expression was observed in 115 dB exposed-animals, progressively returning to baseline levels. This transient elevation may reflect axonal injury triggered by pathological mechanisms happening after NIHL, such as inflammation, oxidative stress, or swellings of the neural tissue. The progressive decrease in SMI312 expression observed 7, 56 and 84d post-exposure may be likely attributed to axonal degeneration events. A transient increase may also represent an early compensatory mechanism aiming to restore input from lower auditory structures (**24,55,56**).

Interestingly, the timing and nature of SMI312 expression pattern differ from previously reported changes in markers like GAP-43 or synaptophysin, which typically emerge weeks or months after exposure (**57–59**). However, GAP-43 or synaptophysin expression is restricted to developing growth cones and has been associated with later phases of synaptic reorganization, where synaptic remodeling and axonal sprouting can occur in order to restore peripheral damage (**57–59**). In contrast, SMI312 targets neurofilament proteins rather than growth cones, being a more sensitive marker to early connectivity changes. Our findings suggest that central neuroplastic responses start immediately after acoustic trauma, in contrast to previous assumptions, and could reflect early signs of appearing hyperactivity in the ACNS following acoustic trauma, which has been linked to tinnitus and hearing loss (**22,23,60**). Moreover, axonal changes in the MGV were more pronounced axonal change than the CIC, possibly reflecting different vulnerabilities between these areas after NIHL. One hypothesis is that the MGV received reduced inhibitory input from the CIC due to early dysfunction of metabolically demanding GABAergic neurons, as proposed by earlier studies of Age-Related Hearing Loss (ARHL) (**61,62**). Large GABAergic cells in the CIC might be first compromised, followed by a loss of GABAergic inhibitory inputs to the MGV (63,64). This loss of inhibitory input could reduce the CIC’s capacity to control excitability in the MGV and inducing higher vulnerability to noise exposure. Likewise, different results were shown by dMRI connectivity, showing only a transient increase in connectivity in 90dB exposed mice 7d post-exposure. This effect may reflect a compensatory mechanism in the MGV to restore peripheral input.

On the other hand, a substantial variability was produced by our image analysis techniques-particularly in FIHC-based quantification –, which may limit the sensitivity to find subtle changes in SMI312 expression in our samples. For this reason, future studies must confirm such detected changes over time. Second, connectivity increases 1d post-exposure were not detected by means of dMRI. Moreover, only transient alterations in dMRI were detected in either the CIC or MGV, which contrasts different studies that described connectivity changes weeks or months after exposure at the MRI level (**41,58,59,65,66**). Differences in specificity and resolution between histological and MRI techniques may explain such discrepancy. In future studies, measurements of structural connectivity using dMRI could be complemented with resting state MRI studies of functional connectivity that may be more sensitive in detecting subtle changes in neuroplasticity after NIHL. Furthermore, differences between studies may arise due to the big spectrum of experimental techniques and experimental models present in the field, as well as different brain regions investigated. NIHL pathological effects are demonstrated to be region-specific, and different noise exposure paradigms or animal models investigated could lead to different conclusions when evaluating the impact of central noise trauma (**6,67**).

Taken together, our study indicates that immediate and region-specific neuroplastic changes occur after NIHL in central auditory pathways. While the intrinsic nature of such changes is still unclear, these early events may be related to long-term alterations in auditory processing, and may represent a critical window for therapeutic, diagnostic or prevention strategies.

### Detection of Neuronal Density Changes

Cell density reductions in the CIC and the MGV after NIHL were not detected by histological or MRI experiments. Absence of GMD changes in VBM is in line with these histological results. Although these findings are in contrast to previous studies by our group (18,19,21), neurodegeneration was probably diminished in the present study despite increased ABR thresholds. In consistence, earlier studies by Aarnisalo et al., 2000 (**30**) and Kurioka et al., 2016 (**68**) found no changes in neuronal density in the VCN, but were able to demonstrate neuropathologies like hair cell damage, apoptosis, reduced VCN volume size and reduced neuronal soma size. Studies by Reuss et al., 2016 (**69**) were not able to quantify neuronal degeneration within the first two weeks after noise exposure in the olivocochlear complex (OC) of the Superior Olivary Complex (SOC). Neurodegeneration is probably more pronounced in other auditory regions not examined in the present study, or pathologies are not attributed to cell and neuron loss. Supporting this hypothesis, human studies detect changes in GMD in canonical and non-canonical ACNS regions of patients suffering from tinnitus, ARHL or occupational NIHL (**46,49–52,70,71**). Since those studies described controversial results, GMD or cell density as a NIHL biomarker needs future research in order to assess its potential therapeutic implications.

At the MRI level, VBM is only a coarse surrogate marker of cell density. One limitation is, that despite the broad use of VBM in the context of neurodegeneration, there is a lack of biophysical models that directly link cell density to GMD. At the histological level, this is the first study in our group to employ FIHC and automated cell counting. These novel methods offer higher specificity of cell detection while removing potential observer bias when performing manual cell counts.

### VNTT and 1H-MRS changes after profound hearing loss

Our study was able to detect minor changes in glutamate and GABA by means of 1H-MRS 84d post-exposure in the 90dB group. Decrease in both Glutamate and GABA after moderate exposure may be attributed to a long-term compensatory mechanism to account moderate hearing damage (**10**). However, VGLUT1, VGLUT2 or VGAT expression levels in the CIC or MGV were not changed significantly in our data after noise exposure. These findings suggest that the observed increases in ABR HTs and/or the early SMI312 expression may not be directly linked to glutamatergic and GABAergic changes. One possible explanation is that that our study lacked sensitivity when trying to estimate changes in Glutamate and GABA concentration. At the MRI level, this is the first study that applied 1H-MRS to evaluate changes in Glutamate and GABA levels *in vivo* after NIHL in an animal model. Additional studies used 1H-MRS to observe changes in Glutamate and GABA, but none of them assessed it specifically after NIHL (**72–76**). Supporting our results, in the CIC, the study by Brozoski et al., 2012 (**72**) detected no significant changes in Glutamate and GABA concentration in an ex vivo animal model of tinnitus. Interestingly, in the MGV, slightly lower GABA concentration in the contralateral site of the exposure, and increased GABA concentration ipsilaterally were found. Studying human patients suffering from tinnitus, studies made by Isler et al., 2022 (**75**) and Sedley et al., 2015 (**76**) detected a reduction in GABA concentration, but within the Auditory Cortex (AC).

To our knowledge, this study is one of the first reports investigating changes in VGLUT1, VGLUT2 or VGAT expression after noise exposure in the CIC and the MGV. Another study by Park et al., 2020 (**77**), showed evaluated the expression of VGLUT1 and VGLUT2 after noise exposure in the CIC using mRNA and protein quantification methods. Supporting the current results, VGLUT1 expression was not affected, but VGLUT2 expression found to be increased 30 days after noise exposure. In the MGV, no studies have investigated the expression of VGLUT1, VGLUT2 and/or VGAT after hearing loss. VNTT expression has been evaluated particularly in the CN after hearing loss, indicating decreased VGLUT1 and increased VGLUT2 expression in this region (**68,77–80**). Observed CN alterations might differ from those in the CIC and MGV, as NIHL presents a prominent region-specific pathology.

Different molecular mechanisms may also play a role in the generation of NIHL central consequences. Several studies found alterations of other vesicular transporters (e.g., VGLUT3), glutamate and GABA receptors (e.g., NMDA, AMPA or GABAAR receptors), synthesis of specific enzymes (e.g., GAD67), or different pathways involved in neurotoxic long-term potentiation events (LTP), throughout the auditory pathway after hearing loss (**54,81–86**). Therefore, future different synaptic targets could be responsible for similar NIHL phenotypes and additional experiments are suggested to assess glutamatergic and GABAergic imbalances in the tissue of interest.

In the present study, one limitation is that, the sensitivity of 1H MRS of glutamate and GABA in our setting requires changes in the mM range. Furthermore, single voxel MRS cannot discriminate the compartment in neural tissue that the neurotransmitter is located in and it is prone to partial volume effects, since an average across a rather large volume is measured. It is thus conceivable, that subtle changes in neurotransmission did not translate to measurable changes in 1H MRS. Similarly, variability produced due to Fluorescence Intensity measurements may had hindered such neurotransmission changes. Hence, further research needs to be done to assess central glutamatergic and GABAergic imbalances.

### Correlation between ABR, MRI and histological parameters

The present study applied a comprehensive multimodal approach to assess relationships among a wide range of biomarkers for NIHL. The selection of biomarkers and analytical methods employed in the study was carried out cautiously according to previous evidences pointing-out their potential involvement in noise-induced plasticity and degeneration. However, it is possible to think that the sensitivity of the selected markers, or the resolution of the applied techniques, was insufficient to detect robust links under our experimental conditions. Nevertheless, as discussed before, many discrepancies arise in the field about how these biomarkers change and which noise exposure models are suitable for reliable studies. Probably, these discrepancies arise because noise-induced neuroplasticity and neurodegeneration is a highly-dynamic and context-dependent process, as multiple compensatory and neurodegenerative pathways may coexist. For instance, early axonal changes detected are transient and probably spatially heterogeneous, escaping detection at the macroscopic level of MRI. Similarly, central synaptic adaptations may not correlate linearly with peripheral damage, due to homeostatic mechanisms or region-specific circuit rewiring. While our results do not support a direct link between the studied parameters, we emphasize the need for refined multimodal and longitudinal approaches to understand the dynamics of auditory plasticity and degeneration. Until then, the diagnostic value of the current biomarkers remains inconclusive.

## CONCLUSION

The current study demonstrates that high-intensity noise exposure triggers rapid but transient neuroplastic changes in central auditory structures during the first moments after noise exposure. Despite persistent elevation of auditory hearing thresholds, no long-term alterations in neuronal density, microstructural MRI markers, or glutamatergic/GABAergic neurotransmission were detected. The absence of significant correlations across functional, histological, and imaging datasets underscores the complexity of behind finding suitable preventive or diagnostic strategies against NIHL. Hence, these findings highlight the need for more sensitive biomarkers to detect and target early CNS consequences of noise-induced hearing loss.

## MATERIAL AND METHODS

### PRE-REGISTRATION

This project was pre-registered in the Open Science Framework (osf.io) (https://osf.io/2aknq/)

### STUDY DESIGN

The study design is illustrated in **Figure 1**. Auditory Brainstem Response (ABR) was used to assess the Hearing Threshold (HT) of the animals. ABRs were performed between one week and one day prior to noise exposure (pre-ABR) to ensure normal hearing. Thereafter, noise exposure was delivered to mice under three distinct paradigms: (1) High noise exposure, for which animals were subjected to a sound pressure level (SPL) of 115 dB for 3h; (2) Moderate noise exposure of 90 dB SPL for 3h; and (3) control sham exposure, for which animals were not exposed to noise under anaesthetic conditions. 1-2 days before MRI another ABR (post-ABR) was performed in order to observe noise-related changes in HT. VBM, dMRI and 1H MRS were performed in different cohorts of mice 1day (1d), 7 days (7d), 56 days (56d) or 84 days (84d) after noise exposure. Immediately after MRI measurements, animals were perfused and brains were carefully dissected. Then, brains were embedded in paraffin blocks for performing FIHC. Then, FIHC stainings were carried out to assess changes in cell density, neurofilament density and vesicular glutamate and GABA transporters. After FIHC, automated and non-automated image analysis techniques were used to analyse both histological and MRI data. Finally, correlation analysis was performed in order to investigate similarities between histology and MRI data.

### ANIMALS

Normal hearing mice (NMRI strain, Charles River, Massachusetts, EUA) of both sexes were used in an adult state (aged 8-10 weeks at noise exposure) when the auditory system is fully developed. Mice did not exceed the age of 6 months during the entire experimental protocol to prevent age-related hearing disorders. Mice were caged in groups of no more than six mice. Each cage included a resting area as well as enrichment elements (plastic tunnels and nesting materials). The animals were kept in a 12/12h dark/light cycle and had constant access to food and water. Between 7 and 8 mice were included in each experimental group included in the present study. The governmental Ethics Commission for Animal Welfare (LaGeSo Berlin, Germany) approved the experimental protocol (approval number: G0256/19). The study protocol was in accordance to the EU Directive 2010/63/EU on the protection of animals used for scientific purposes.

### NOISE EXPOSURE

The chosen noise exposure paradigm was based on earlier studies on neuronal degeneration and cell death mechanism detection after noise exposure in order to ensure similar experimental conditions, taking into account the hearing range of mice (≥1Hz and ≤100 kHz) (Basta et al., 2005; Gröschel et al., 2010, 2024). The noise was delivered binaurally inside a sound-proof chamber (80 cm × 80 cm × 80 cm, minimal attenuation 60 dB) by a speaker (HTC 11.19; Visaton, Haan, Germany) placed right above the animal’s head and connected to an audio amplifier (Tangent AMP-50; Aulum, Denmark). dB SPL levels were calibrated using a sound level meter (Voltcraft 329; Conrad Electronic, Hirschau, Germany) placed at the animal’s head position. Temperature inside the chamber was measured with a temperature controller (GTH 1200A Digital thermometer, Greisinger Electronic GmbH, Germany), and a heating pad (Thermolux Wärmeunterlagen, Witte & Sutor GmbH, Germany) was used to ensure constant body temperature (36 °C) during the experiments. Noise exposure experiments were conducted under anesthesia (4 mg/kg xylazine and 100 mg/kg ketamine). Animals were selected randomly to undergo noise exposure treatment (115 dB or 90 dB SPL) for 3h to broadband white noise (5-20 kHz). A separate group of animals was used as the sham control (Ctrl) group, and did not receive any noise exposure treatment. The control animals were equally anesthetized and kept in the heating chamber under similar conditions. To ensure constant anesthetic state in the mice during noise exposure, 30% of the initial anesthesia dose was injected by a subcutaneous catheter approximately every 30 minutes during noise exposure treatment. Mice were constantly monitored using a video camera (Logilink UA0072A, Logitech, Lausanne, Switzerland) placed inside the soundproof chamber. After noise exposure, animals were placed in their home cages and kept in the animal facility with continuous access to food and water until subsequent measurements were performed.

### AUDITORY BRAINSTREM RESPONSE (ABR)

ABR recordings were performed one-week prior to noise exposure (pre-ABR) under anesthesia (100 mg/kg ketamine, 4 mg/kg xylazine) to assess the HT before noise exposure. 1-2 days prior to MRI measurements, ABR recordings (post-ABR) were repeated to detect changes in HT after noise delivery. ABR measurements were performed inside the soundproof chamber under temperature and anesthetic conditions as used during noise exposure. Subdermal needle electrodes (Neuro.Dart SD57-426-M, Spes Medica Inc., Italy) were positioned on the forehead (reference, green), mastoid (recording, red), and at one foot (ground, black). In order to prevent electrical interference, a metal cage was positioned above the animals. Tones were presented binaurally at various sound pressure levels (SPL) using a high-frequency loudspeaker (HTC 11.19; Visaton, Germany) placed right above the animal’s head. The presented tones were adjusted using a sine-wave generator (FG 250 D, H-Tronic, Germany) connected to an audio amplifier (Yamaha A-S 201, Yamaha Corp., Japan). ABR waveforms were recorded by the MC_Rack software (MC_Rack, Multichannel Systems, Harvard Bioscience Inc.) and a recording amplifier (USB-ME16-FAI-System, Multi-Channel Systems, Germany). In the present study, ABR specific responses were recorded at four different frequencies (4, 8, 16, and 32 kHz) during both pre-ABRs and post-ABRs. After pre-ABRs and post-ABR, mice were returned to their home cages and kept in the animal facility until experimental day. Following post-ABRs, the individual hearing threshold shift was calculated for each tested frequency. Results were averaged for each experimental group and experimental time point, and mean threshold shifts (Mean (dB) ± Standard Deviation) were calculated and compared to control animals. Post-ABR recordings were not performed in the 1d post-exposure group as previous studies from our group showed difficulties to detect any ABR response 1d after acute noise (54), as well as to minimize the dropout rate of our experimental subjects.

### MAGNETIC RESONANCE IMAGING (MRI)

MRI was performed using a 7 T animal scanner (BioSpec 70/20 USR, Bruker, Ettlingen, Germany) with a cryogenically cooled transmit/receive mouse head surface coil and Paravision 6.0.1 software. Measurements started 1-2 days after post-ABR measurement. Animals were continuously anesthetized using Isoflurane (1.5%) in a 70%/30% N2O/O2 mixture. An MR-compatible monitoring system (SA Instruments, Stony Brook, NY, USA) was used to continuously control animal temperature via a rectal probe and respiration rate via a pressure-sensitive pad under the thorax. Animals were placed on an animal holder inside the MRI with ear and tooth bar fixation to avoid movement and in prone position. The protocol consisted of T2-weighted (T2w) MRI for VBM of gray matter changes, dMRI for connectome reconstruction, and 1H MR spectroscopy (1H MRS) for neurotransmitter quantification.

### VOXEL-BASED MORPHOMETRY (VBM)

T2w images were acquired with a 2D-RARE with repetition time (TR) = 5300 ms, effective echo time (TE) = 33 ms, echo spacing (TE) = 11 ms, RARE factor = 8, 2 averages, 50 contiguous axial slices with a slice thickness of 0.32 mm, field of view (FOV) = 19.2 × 19.2 mm^2^, matrix size MTX = 256 × 256, bandwidth BW = 34722 Hz, 3 averages and total acquisition time TA = 8:29 min. Images were segmented into tissue probability maps of gray/white matter and cerebrospinal fluid and nonlinearly registered to a custom brain atlas with 38 anatomical regions of the mouse auditory system (19 regions per hemisphere) derived from the Allen mouse brain atlas (CCFv3) in ANTx2 (https://github.com/ChariteExpMri/antx2). Using the atlas, modulated (i.e. tissue probability multiplied with jacobian determinant of the registration) mean gray and white matter density (GMD, WMD) in CIC and MGV were measured for further group statistical analyses. For all readouts reported in this study, left and right CIC/MGV were merged, i.e. analyzed as one single region of interest.

### DIFFUSION MRI (dMRI)

dMRI images were acquired with a multishot 2D spin echo EPI sequence with geometry matching the T2w images but lower resolution (MTX=192 x 192, 40 contiguous slices, slice thickness 0.4 mm, 6 EPI segments, TR=2200 ms, TE=32.4 ms, double sampling on, BW=300 kHz, diffusion gradient duration/separation= 3.9 ms/16 ms) and multishell diffusion encoding (5 b=0 images, 6 directions with b=100 s/mm2, 13 directions with b=900 s/mm2, 25 directions with b=1600 s/mm2, 37 directions with b=2500 s/mm2, TA=18:42 min). Diffusion directions were calculated with the online tool available at http://www.emmanuelcaruyer.com/q-space-sampling.php (**87**). The number of directions varied linearly with diffusion wave vector q (b∼q2), i.e., with a square root dependency N(b)∼b^1/2^. The auditory brain system atlas was registered to the T2-weighted images using the inverse transform calculated for VBM and further registered with an affine transform (12 degrees of freedom) to the diffusion MR images using ANTx2 (https://github.com/ChariteExpMri/antx2). Diffusion MR images were processed in mrtrix (https://www.mrtrix.org) and custom tools based on network analysis as described previously (**88,89**). Scripts are available via github (https://github.com/ChariteExpMri/rodentDtiConnectomics). The processing pipeline consisted of: (1) Conversion of Bruker raw data into NIFTI image format; (2) Denoising, Gibbs ringing removal, bias field correction, eddy current correction and motion correction; (3) Calculation of conventional diffusion tensor imaging maps of axial and radial diffusivity and apparent diffusion coefficient (ad, rd, adc) and of fractional anisotropy (fa); (4) Diffusion orientation function reconstruction using constrained spherical deconvolution; (5) Connectome reconstruction using streamline tractography and SIFT2 optimization; (6) Reconstruction of connectivity matrices counting the number of streamlines from atlas region to region. Mean ad, rd, adc and fa within CIC and MGV and the sum of terminating streamlines originating from each of those regions were derived from the connectivity matrices.

### PROTON MAGNETIC RESONANCE SPECTROSCOPY (1H-MRS)

Single voxel 1H MRS was performed using a stimulated echo acquisition mode (STEAM) sequence following local shimming (MAPSHIM) across a single cuboid voxel placed in the target region (CIC or MGV) with TR/TE/mixing time/=2500 ms/3 ms/ 10 ms, VAPOR water suppression, outer volume suppression, acquisition duration 620.41 ms, 2048 points, bandwidth 3301.06 Hz, dwell time 151.47 µs, 0.81 Hz/point, number of averages 600, TA=25:00 min. Voxel size was 1.0 mm (lateral) x 1.4 mm (ventral-dorsal) x 1.2 mm (rostral-caudal) for CIC and 0.8 mm x 1.2 mm x 1.4 mm for MGV. A water-unsuppressed scan was acquired for water-scaling. Spectra were fitted using LCModel (http://s-provencher.com/lcmodel.shtml), and water-scaling was used to calculate absolute metabolite concentrations of Glu and GABA.

### FLUORESCENCE IMMUNOHISTOCHEMISTRY (FIHC)

Immediately after MRI measurements, mice were perfused via the left heart chamber first with NaCl solution (0.9%) and secondly with 4% paraformaldehyde (Sigma, Germany). Brains were carefully removed from the skull and preserved in 0.1% PFA. Neural tissue was post-fixed in formalin for 48h. Brains were embedded in paraffin blocks using an embedding workstation (Epredia HistoStar Embedding Workstation, Massachusetts, Thermo Fisher Scientific), and 5-µm-thick slices were cut in the frontal plane using a rotation microtome (Rotationsmikrotom Microm HM 325, Thermo Fisher Scientific). Sectioning was carried out from the beginning of the Cochlear Nucleus (CN) (Bregma, –6.48 mm; Interaural, –2.68 mm) until the end of the MGV (Bregma, –2.80 mm; Interaural, 1.00 mm), following the histological brain atlas of Paxinos and Franklin (**90**). After sectioning, brain slices were first embedded in Roti Histol organic solvent (Roti Histol, Carl Roth, Karlsruhe, Germany) to remove paraffin (10 min., RT). Subsequently, slices were rehydrated using descending alcohol concentrations of 90% (1x, 5 min, RT), 80% (1x, 5 min, RT) and 70% (1x, 5 min, RT), and finally washed in PBS 1X (Gibco, New York, USA) (1x, 5 min., RT). Sections were incubated in 10 mM HIER Citrate Buffer (Biozol, Germany) for performing antigen retrieval steps through Heat Induced Epitope Retrieval Immunofluorescence protocol (HIER-IFA). HIER-IFA protocol was carried out by cooking brain slices in HIER Citrate Buffer inside a high-pressure cooker (Pressure Cooker, RC-HPC6L, Royal Catering, Poland), filled with 500 ml of dest. water (dH_2_O) for 5 minutes. Then, sections were cooled down for 30 minutes on running tap water, and washed 5 minutes in PBS 1X. Afterwards, the tissue of interest was delineated with hydrophobic liquid barrier (RotiLiquid, Carl Roth, Germany) and blocked in 5% normal goat serum (NGS), 0.3% Triton and PBS 1X for 30 minutes at RT. Thereafter, depending on the experiment, different primary antibodies were used. For NeuN labelling, the primary antibody NeuN (D3S3I) Rabbit mAb primary anti-NeuN antibody (Cell Signaling Technology, Massachusetts, USA) diluted 1:100 in 1% NGS, 0.3% Triton and PBS 1X was used. For NF labelling, primary antibody incubation solution was prepared using purified mouse SMI312 anti-neurofilament marker antibody (pan axonal, cocktail) (Biolegend, San Diego, USA) diluted 1:1000 in 1% NGS, 0.3% Triton and PBS 1X incubation solution. For VNTT labelling, three different primary antibodies were used: polyclonal chicken antibody SySy 135 316 against VGLUT1 (Synaptic Systems, Inc.); polyclonal guinea pig antibody SySy 135 404 against VGLUT2 (Synaptic Systems, Inc.), and polyclonal rabbit antibody SySy 131 002 against VGAT (Synaptic Systems, Inc.) diluted 1:500 in 1% NGS and 0.3% Triton and PBS 1X incubation solution. All primary antibody incubation steps were performed during 24h at 4°C inside a humid chamber to avoid tissue drying. On the second day, slices were washed in PBS 1X three times during 10 minutes in agitation. Following, incubation of secondary antibodies was performed. For NeuN, we used Alexa Fluor 488 Goat anti-Rabbit IgG (H+L) (Thermo Fisher Scientific, USA) diluted 1:250 in 1% NGS, 0.3% Triton and PBS 1X incubation. For NF labelling, we used Alexa Fluor 488 Invitrogen Goat anti-mouse IgG (H+L) (Thermo Fisher Scientific, USA) diluted 1:250 in 1% NGS, 0.3% Triton and PBS 1X incubation. For VNTT labelling, we incubated simultaneously three different secondary antibodies diluted 1:500 in 1% NGS, 0.3% Triton and PBS 1X: For VGAT labelling, we used Alexa Fluor 488 Goat anti-Rabbit IgG (H+L) (Thermo Fisher Scientific, USA); For VGLUT1 labelling, we used Alexa Fluor 647 Invitrogen Goat anti-chicken IgY (H+L) (Thermo Fisher Scientific, USA); For VLGUT2 labelling, Alexa Fluor 546 Invitrogen Goat anti-Guinea Pig IgG (H+L) (Thermo Fisher Scientific, USA) was used. Secondary antibodies were incubated during 1h at RT in dark. Finally, sections were rinsed three times in PBS 1X (10 min, RT, dark). Subsequently, each microscope slide was incubated with 300 µl of DAPI 300 µM in PBS 1X, 5 min at RT in dark conditions (4’,6-Diamidino-2-phenylindole, Dihydrochloride, D1306, Thermo Fisher Scientific, USA). Negative control (NC) stainings were performed for obtaining proper background fluorescence intensity. In NC stainings, secondary antibodies as well as DAPI incubation solution were added only, without primary antibody incubation. Stained microscope slides were mounted in Roti Mount FluorCare mounting media (RotiMount FluorCare, Carl Roth GmbH & Co), covered with coverslips (Deckgläser 24×60, H878, Carl Roth GmbH & Co.), sealed with nail polish and placed horizontally O/N at 4 °C inside microscope slide boxes. Sealed slides were kept at 4 °C in dark conditions until image acquisition.

### IMAGE ACQUISITION: HISTOLOGY

For acquisition of histological images of the CIC and MGV an inverted widefield fluorescence microscope (Ti2, Nikon, Japan) controlled by NIS-Elements (Nikon) was used. A LED system (SpectraX, Lumencor, USA) provided the fluorescence excitation (ex). The emission (em) was detected by a sCMOS camera (pco.edge 4.2 USB, Excelitas pco, Germany). The system provided the following fluorescence channels (numbers in nm): DAPI (ex 395/25, em empty), GFP (ex 475/28, em 519/26); YFP (ex 511/16, em 540/30); Cy5 (ex 635/22, em 697/60) in combination with the following multi-band dichroic mirrors: DAPI/GFP/mCherry/Cy5 (quad ET435/33, 526/20, 595/38, 695/63), CFP/YFP (quad ET475/25, 537/30, 644/92, 806/100), FRET (455LP). NeuN and DAPI stained slices were imaged using the 20x objective (Plan Apo Ph2 DM, NA 0.75, WD 1000 µm). SMI312 (image size 336.16 x 335.20 µm) and VGAT, VGLUT1, VGLUT2 stained slices (image size 336.16 x 335.20 µm) were imaged using the 40x objective (Plan Apo (Air), NA 0.95, WD 250-170 µm). Images from both positive and negative control staining were acquired under the same settings (LED power, exposure time) in order to quantitatively compare the fluorescence intensity values. Up to 12 images from both hemispheres were acquired for the CIC and MGV and for each experimental subject included in the study. Imaging was performed at the Advanced Medical BIOimaging Core Facility of the Charité-Universitätsmedizin Berlin (AMBIO).

### IMAGE ANALYSIS: HISTOLOGY

For histological data, image analysis was performed using automated Fiji/ImageJ (NIH, USA) macros developed in collaboration with AMBIO. Automated cell counting was performed to assess total cell (DAPI⁺) and neuronal (NeuN⁺/DAPI⁺) densities. Fluorescence signals were segmented using Cellpose (https://www.cellpose.org) in a Python environment, with image intensities normalized between the 1st and 99th percentiles. The ‘nuclei’ model was used for DAPI⁺ segmentation and the ‘cyto’ model for NeuN⁺. Total cells (DAPI⁺) were counted from the DAPI channel, while neurons (NeuN⁺/DAPI⁺) were identified by overlapping fluorescent signals from the DAPI and NeuN channels. All processing steps were implemented in a custom Python Jupyter Notebook. Results are given in DAPI^+^ or NeuN^+^/DAPI^+^ counts / area (Mean ± Standard Deviation (SD)).

Relative protein expression levels of SMI312, VGLUT1, VGLUT2, and VGAT were quantified using a custom Fiji/ImageJ (NIH, USA) macro. Images were acquired with the field of view (FOV) size kept constant for both positive control (PC) and negative control (NC) samples. Neurons were segmented via histogram-based thresholding (Yen’s method; https://doi.org/10.1109/83.366472) following ImageJ’s ‘rolling ball’ background subtraction. The following formula was applied to retrieve the ‘Corrected Total Cell Fluorescence’ (CTCF) by subtracting the fluorescence intensity of the respective control group from each measurement.

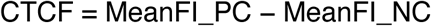

MeanFI_PC refers to fluorescence intensity values measured from PC samples, while MeanFI_NC refers to values from NC samples. NC samples were derived from negatively stained slices for each staining session and auditory brain region to minimize methodological and biological variability in fluorescence intensity. Results are given in Mean Fluorescence Intensity (SMI312_CTCF_, VGLUT1_CTCF_, VGLUT2_CTCF and_ VGAT_CTCF_) levels (Mean ± SD).

### CORRELATION ANALYSIS

In order to evaluate potential relationships between different parameters investigated, a correlation analysis was performed. Spearman correlation ranks between ABR, histological and MRI-derived variables were calculated. Analysis was performed without discriminating between timepoints and noise exposure conditions, using both SPSS (version 29.0.0.0. (241), IBM SPSS Statistics) and GraphPad Prism 9.4.1 software.

### STATISTICS

This study has a mixed design. A between-subject design with 4 groups (experimental time point: 1d, 7d, 56d, 84d) for 3 treatments (115 dB, 90 dB, and Ctrl), with 3 within-subject levels was established. Each of the hypotheses were tested separately for each auditory CNS area of interest (CIC and MGV). Since the primary objective of the current study was to track the pathological effects of various noise exposure conditions at 4 time points, the resulting data was compared between the experimental groups or treatments (115 dB, 90 dB and Ctrl) for each brain region within each experimental time point (1d, 7d, 56d and 84d). Data is always shown in mean ± Standard Deviation (SD). The resulting means of the experimental groups were compared using one-way ANOVA and the Tukey post-hoc test was applied to account for multiple comparisons, when homogeneity of variances in the data was assumed. In cases where homogeneity of variances was not given, Welch’s ANOVA followed by the Games-Howell post-hoc multiple comparisons test was performed. Taking into account the sample size collected and the robustness of the previously described statistical analysis against the normality assumption, violations of normality in data distribution were not taken into account in the present study. During the correlation analysis, Spearman correlations ranks were calculated due to the lack of normality and linear relationships between investigated variables. The software SPSS (IBM SPSS Statistics, version 29.0.0.0. (241), New York, USA), Microsoft Excel and GraphPad Prism 9 software were used for conducting statistical analyses. The alpha error level was set at p ≤ 0.05.

## ACKNOWLEDGEMENTS

Authors recognize the support of Dr. Max Meuser and Dr. Lenneke Kiefer from the Center for Medicine Technology of the Unfallkrankenhaus Berlin due to his valuable support during the discussion and the execution of the current experiments. We would also like to thank Dr. Anja Kühl from the Institute of Pathology of the Charité Universitätsmedizin Berlin, who gave key important advises in order to refine the current fluorescence immunohistochemistry protocols. We also thank the Advanced Medical BIOimaging Core Facility of the Charité-Universitätsmedizin Berlin (AMBIO) for support in acquisition and analysis of the imaging data.

## Figure Legends

**Supplementary Figure 1.** Microstructural connectivity changes after noise exposure. Diffusion MRI (dMRI) parameters were assessed in the central inferior **colliculus (CIC; left panels, A–A″)** and the ventral medial geniculate body **(MGV; right panels, B–B″)** at different time points following noise exposure. Group comparisons revealed no statistically significant changes in axial diffusivity **(AD; A, B)**, apparent diffusion coefficient **(ADC; A′, B′)**, or radial diffusivity **(RD; A″, B″)** in either region across conditions and time points (p > 0.05 for all comparisons). Data is presented as mean ± standard deviation (ns = not significant, ***: p < 0.05, **: p < 0.01, ***: p < 0.001).

