## Supplementary figures and images for "Temporal dynamics of neuroplasticity and neurodegeneration in the central auditory system following noise-induced hearing loss: A multimodal imaging and histological study"

### Central neurodegeneration and neuroplasticity after NIHL_Supplemental_100925.pdf

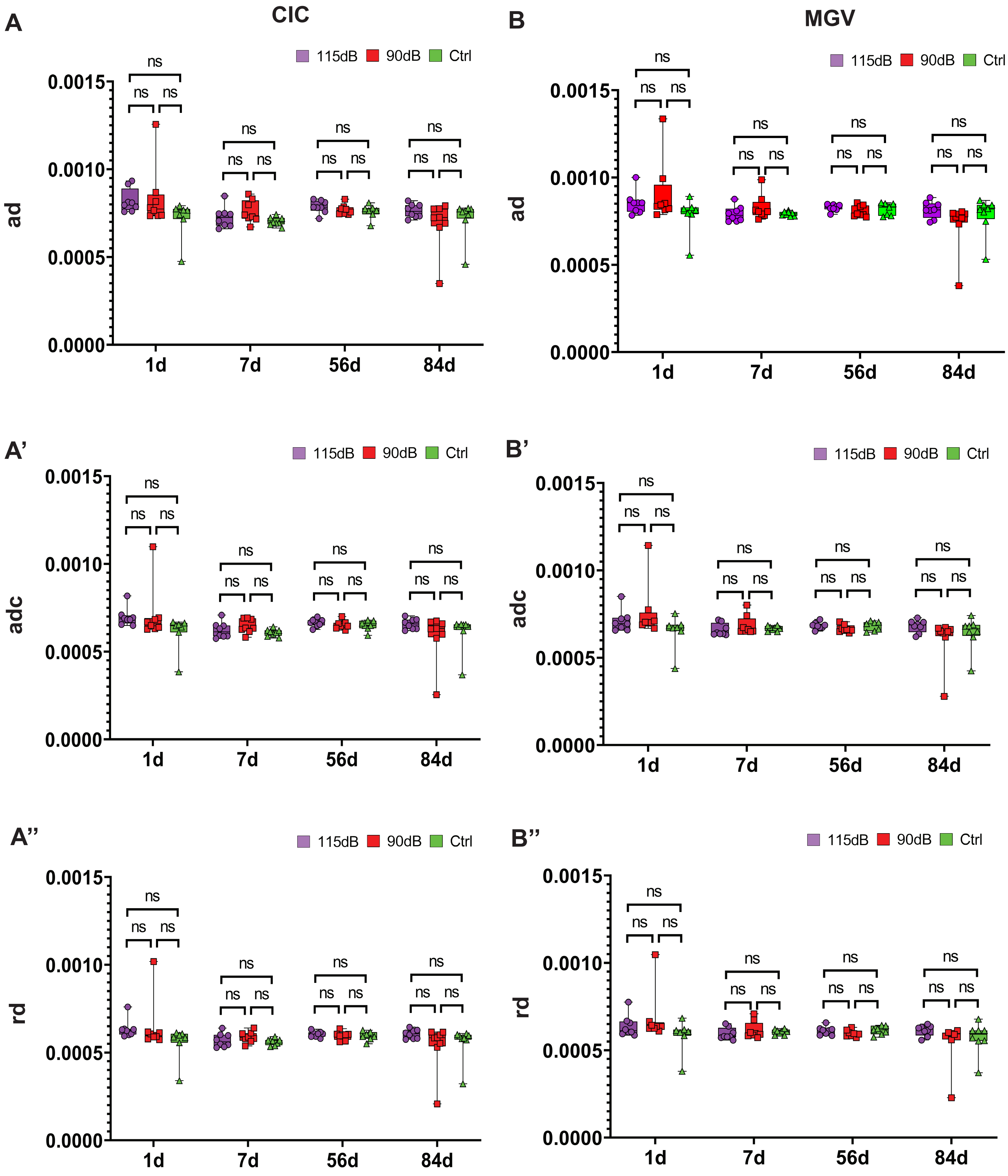


**Suplementary Figure 1**
